# Inferring the genetic architecture of expression variation from replicated high throughput allele-specific expression experiments

**DOI:** 10.1101/699074

**Authors:** Xinwen Zhang, J.J. Emerson

**Affiliations:** Department of Ecology and Evolutionary Biology, University of California Irvine, Irvine, CA 92697

## Abstract

Gene expression variation between alleles in a diploid cell is mediated by variation in *cis* regulatory sequences, which usually refers to the differences in DNA sequence between two alleles near the gene of interest. Expression differences caused by *cis* variation has been estimated by the ratio of the expression level of the two alleles under a binomial model. However, the binomial model underestimates the variance among replicated experiments resulting in the exaggerated statistical significance of estimated *cis* effects and thus many false discoveries of *cis*-affected genes. Here we describe a beta-binomial model that estimates the *cis*-effect for each gene while permitting overdispersion of variance among replicates. We demonstrated with simulated null data (data without true *cis*-effect) that the new model fits the true distribution better, resulting in approximately 5% false positive rate under 5% significance level in all null datasets, considerably better than the 6%-40% false positive rate of the binomial model. Additional replicates increase the performance of the beta-binomial model but not of the binomial model. We also collected new allele-specific expression data from an experiment comprised of 20 replicates of a yeast hybrid (YPS128/RM11-1a). We eliminated the mapping bias problem with *de novo* assemblies of the two parental genomes. By applying the beta-binomial model to this dataset, we found that *cis* effects are ubiquitous, affecting around 70% of genes. However, most of these changes are small in magnitude. The high number of replicates enabled us a better approximation of *cis* landscape within species and also provides a resource for future exploration for better models.

## Introduction

Variation in gene expression contributes significantly to phenotypic variation (Jacob and Monod 1961; Mcclintock 1956). Consequently, gene regulatory elements have long been thought to be an important target of natural selection comparable in significance to variation in the proteome (Ohno 1972; King and Wilson 1975; Wray 2007). The genetic architecture of variation in gene regulation can be decomposed into *cis* variation and *trans* variation. The *cis* variation affects expression differences between two individuals in a non-diffusible manner (e.g., a mutation on a promoter region), while *trans* variation affects the expression difference in a diffusible manner (e.g., a coding region mutation on a transcription factor) (Emerson and Li 2010; Wittkopp, Haerum, and Clark 2008).

In an F1 hybrid cell, two alleles of the same gene are exposed to the same diffusible elements, so any difference between the alleles’ expression must be encoded by features linked to the gene itself (i.e., *cis* variation). By measuring the allele-specific expression of all genes in hybrid cells, we can measure the magnitude of cis variation (*cis*-effect) and detect *cis*-affected genes (Signor and Nuzhdin 2018; Emerson and Li 2010). The *cis* effect parameter (e_cis_) for a gene is defined as the ratio of the expression from allele 1 and allele 2 (Emerson et al. 2010; Schaefke et al. 2013). However, previous allele-specific expression studies using RNA-seq for *cis*-effect typically employed 1-3 hybrid replicates in binomial framework (Emerson et al. 2010; Schaefke et al. 2013; Metzger, Wittkopp, and Coolon 2017; Rhoné et al. 2017; Mack, Campbell, and Nachman 2016; McManus et al. 2014; Bell et al. 2013), which assumes that the read counts for each allele among replicates can be modeled as a Poisson random variable.

The actual variance among RNA-seq experiments is known to be overdispersed, and consequently, the single Poisson parameter is inadequate to model both the mean and variance. The negative binomial distribution instead has been shown to fit better than Poisson in many differential expression studies (Robinson and Smyth 2007; Schurch et al. 2016; Gierliński et al. 2015). The negative binomial distribution is equivalent to the compound gamma-Poisson distribution, where the lambda parameter of Poisson is a gamma-distributed random variable. The two parameters of the negative binomial permit the mean and variance to vary independently. Therefore, we modeled allelic expression for each gene with a negative binomial distribution instead of a Poisson distribution. Under this assumption, the cis-effect e_cis_ is beta-binomially distributed with an overdispersion parameter compared with the binomial distribution (See Materials and Methods: *Cis* variation estimation).

We compared the false positive rates of the two models with simulated null datasets where no true *cis* effects exist. We found that the binomial model has high false positive rate even with a large number of replicates, but the beta-binomial model improves with increased replication, attaining a 5% false positive rate as expected.

We also grew 20 replicates of hybrid from the cross of yeast *Saccharomyces cerevisiae* strains YPS128 and RM11-1a to estimate *cis* variation. YPS128 is a woodland stain (Sniegowski, Dombrowski, and Fingerman 2002) and RM11-1a is a derivative of a vineyard strain (Brem et al. 2002). We used RNA-seq for allele-specific counts and estimated the gene-wise e_cis_ with both models. In terms of power, both models improve as replication increases. We found from this experimental data that ~70% of the total 4710 informative genes have a significant cis difference. Around 2% of the total genes have a greater than 2-fold difference significantly.

Estimated from the simulated null data, 20% - 30% genes lacking a true *cis* effect would be falsely classified as significant by the binomial model. In our experimental data, the beta-binomial model and binomial model differ by ~5% in the number of significant *cis* affected genes (Figure 4), which is less than the 15% - 25% difference in false-positive rate estimated from the null data. This could perhaps be explained by the possibility that the two strains are sufficiently diverse that most of the genes are true positives. However, for closely related species (or strains) with less differential gene expression, a 5% false positive rate would contribute a much higher proportion to the total number of differentially expressed genes.

This allele-specific study demonstrated the advantage of the beta-binomial model over the binomial model and the salutary effect of using high replication. The high number of replicates of hybrid samples between the two yeast strains enabled us a better approximation of *cis* landscape within species. It also provides a resource for future exploration of better models.

## Materials and Methods

### Yeast strains and preparation of hybrid samples

To prepare the hybrid strain, we mixed a single colony of YPS128 strain (MAT ɑ; ura3::kanMX; HO::HygMX; lys2::ura3) and a single colony of RM11-1a strain (MATa; leu2Δ0; ura3Δ0; HO::kanMX) together in 100 ul YPAD, put the mix in 30 °C for 4 hours, then we poured 50ul of mixed cells into a dropout plate (-leu, -lys), and struck to get one single diploid colony.

We picked one single diploid colony and struck it on the standard YPAD plate for hybrid sample collection. We then collected 20 independent hybrid samples started from this YPAD plate. Each sample was generated by the following procedure:

One single colony was taken from the YPAD plate and was cultured overnight. It was then diluted to OD 0.05 in 5ml YPAD and grow until OD 0.7-0.8 in 30 °C with 220 rpm shaking. The yeast culture was then distributed in Eppendorf tubes by 1 ml per tube, centrifuged with 9000 rpm to remove the supernatant, snap-frozen in liquid Nitrogen and finally stored at −80°C for DNA and RNA extraction.

### DNA extraction and sequencing

We extracted the DNA of these 20 hybrid samples using the Yeast DNA Extraction Kit (Thermo Scientific 78870). After extraction, we used the Nextera DNA Library Preparation Kit (Illumina) to make 20 libraries with unique barcode combination (Nextera Index Kit) and pooled them together before sequencing. We sequenced the pooled library in UC Davis Genome Center (http://dnatech.genomecenter.ucdavis.edu/) with 1 Lane of mid-output Nextseq PE75. We then demultiplexed (Renaud et al. 2015) the pooled reads and got a total of 95.3 million reads for the 20 replicates.

### RNA extraction and sequencing

We extracted the RNA of these 20 hybrid samples using the TRIzol Plus RNA Purification Kit (Invitrogen). Transcriptome libraries were made by the Smart-seq2 protocol (Picelli et al. 2014). The 20 Libraries were pooled together and sequenced in UC Davis Genome Center with 4 Lanes of high-output Nextseq PE75. After demultiplexing (Renaud et al. 2015), we got a total of 1530.8 million reads for the 20 replicates.

### Sequencing and assembly of YPS128 and RM11-1a genome

We assembled our YPS128 and RM11-1a genome and used them as the reference genomes in mapping DNA/RNA reads. We extracted whole genome DNA of YPS128 strain and RM11-1a strain using the QIAamp DNA Mini Preparation Kit (Qiagen), prepared Nextera DNA library (Illumina) and sequenced the pooled library with 1 Lane of Miseq PE75, which generated an 88X coverage for the YPS128 strain and a 102X coverage for the RM11-1a strain. We also generated long DNA reads with Oxford Nanopore (Rapid sequencing) for RM11-1a strain and got a 59X coverage. Since our Nanopore experiment failed for the YPS128 strain, we downloaded its Pacbio long reads from this project (Yue et al. 2017) which gives a 230X coverage.

For YPS128 strain, we used Dextractor (https://github.com/thegenemyers/DEXTRACTOR) to extract fastq sequences from the original h5 files. Then we used Canu (Koren et al. 2017) for raw assembly and finisherSC (Lam et al. 2015) for gap fixing, followed by two rounds of quiver (https://github.com/PacificBiosciences/GenomicConsensus) correction. We further polished the assembly with Illumina short reads using pilon (Walker et al. 2014) and pacbio long reads again using quiver followed by one final round of pilon. We ended up with an assembly with NG50=808.6K and Busco score (Simão et al. 2015) of 94.4%(fungi).

For RM11-1a strain, we used Albacore (ONT software version 2.2.7) for nanopore long reads base-calling. Then we used Canu (Koren et al. 2017) for raw assembly followed by finisherSC (Lam et al. 2015) for gap fixing, then corrected the raw assembly by three rounds of Racon (https://github.com/isovic/racon). We further polished the assembly with the Illumina short reads using Pilon (Walker et al. 2014) and nanopore long reads again using Racon. We did the pilon-racon for two rounds and wrapped up with four rounds of pilon. Finally, we obtained an assembly with NG50=919.8K and Busco score (Simão et al. 2015) of 93.7% (fungi).

The qualities of the assemblies are further evaluated with QV estimation. We aligned the Illumina reads used for polishing to the final assembly using bwa mem (Li and Durbin 2009) with default parameters. Following (Koren et al. 2018), we used freebayes (V.1.2.0-4) (Garrison and Marth 2012) to estimate the number of SNPs and indel variants with the command “freebayes -C 2 -0 - O -q 20 -z 0.10 -E 0 -X -u --ploidy 1 -F 0.75 -f asm.fasta asm_nodup.bam > asm.vcf”. Total based changed E (inserted, deleted, substituted) was summed and divided by the total number of bases (T) with minimum coverage 3. QV was calculated as −10log10(E/T).

### Collect DNA/RNA read counts

#### Identify variants between YPS128 and RM11-1a

The reads from hybrid samples are unidentifiable of which parental genotype they belong to if they don’t overlap with any variant (SNPs or Indels) between the two parental strains. So we first extracted a list of SNPs and Indels by comparing the YPS128 assembly and RM11-1a assembly using MUMmer((Kurtz et al. 2004) MUMmer/3.23: nucmer; show-snps). For conservativeness, we did it in both directions (using YPS128 as query, RM11-1a as subject and then exchange) and only retained the SNPs and Indels that appear in both comparisons.

#### Mapping DNA reads with two references

We next mapped the DNA reads of the 20 hybrid samples to both assemblies using bowtie2 ((Langmead and Salzberg 2012) bowtie2.2.7) and got 40 mapping files. We then counted the allele-specific number of reads hitting each variant position with Samtools ((Li et al. 2009) Samtools 1.9: mpileup setting -q to 5 to ignore multi-hits reads) and customized scripts (count_pileup.py: count the number of reads mapping to the reference allele and alternative allele respectively using mpileup output file as input).

We found that the mapping always biases towards the reference genome. In hybrid DNA samples, the reads from YPS128 genome is expected to be of the same amount as from RM11-1a genome. However, when YPS128 assembly was used as the reference genome, the sum of reads assigned to YPS128 allele across all variants is around 1.4 fold more than the sum of reads assigned to RM11-1a allele in all of the 20 hybrids. This also happened when RM11-1a was used as the reference genome. The sum of reads assigned to RM11-1a allele across all variants is around 1.4 fold more than the number of reads assigned to YPS128 allele (Figure S1).

One main reason for this mapping bias is that when one assembly was chosen as the reference genome, the reads from the alternative genome in the hybrid sample are not as likely to map to the correct genomic position because of the variant. So we conceived that the alternative counts for each SNP/indel in the mapping results are underestimated while the reference counts are more reliable. Thus, we only kept the YPS128 allelic reads from mapping results using YPS128 as the reference genome and RM11-1a allelic reads from mapping results using RM11-1a as the reference genome. Some variant positions are close to each other and the reads that cover both of them would be counted repeatedly when summing up the counts, so we also unioned the reads from each allele using the reads’ names as identifiers. After this operation, we reduced most of the mapping bias, but the total read counts still biased towards RM11-1a genome by around 1% (Figure S2).

#### Identifying suspected loci causing mapping bias

Another possible source for mapping bias are the errors in genome assembly and the incoordination between the assembly and the real genotype in hybrid (the YPS128 strain’s Pacbio long-reads used in assembly is not from this project), or regions that the sequencing probability for two alleles is extremely different. In these loci, nearly all the allelic read counts would be assigned to one of the genomes. As these kinds of loci accumulate, the bias would be reflected in the total read counts. Thus, we check the reads that cover each variant position to see whether the nucleotide information provided by the short reads in hybrid samples match with the variant we got from genome comparison. For example, If the SNP pair is A on YPS128 and C on RM11- 1a from the comparison of assemblies, short reads with A and short reads with C on the corresponding positions are both required to exist in all mapping results. Variants without sufficient short reads support were removed for downstream analysis (12793 positions are removed from total 82029 positions in YPS128; 12326 positions are removed from total 81574 positions in RM11-1a). After the removal of those positions, we recounted the read counts overlapping with the remaining positions and also unioned the reads covering consecutive positions (group_reads.py, yps5rmB_gc.py), the mapping bias was then sufficient small to be ignored (Figure S3).

#### Mapping expression reads and collecting allele-specific read counts

We first annotated the two assemblies with CrossMap ((Zhao et al. 2014) v0.2.8: using S.cer reference annotation), and label each variant position with gene name. The variant positions that are not in any gene regions or overlap with two gene range (Some gene overlaps in yeast) are further removed. There are 37487 variant positions retained, which cover 4710 genes.

We then mapped the Expression reads using bowtie2 (bowtie 2.2.7: there are very limited intron regions in the yeast genome, so we didn’t choose an RNA splice-sites aware mapping tool) to both two assemblies. Same as the procedure for DNA reads counting, we collected counts from only reference allele for each retained position.

Finally, we aggregated the read counts of variant positions under the same gene name and counted allelic reads with samtools and customized scripts as we did for DNA counts (group_reads.py, yps5rmB_gc.py). The total read counts of YPS128 allele and RM11-1a allele are almost the same in the 20 hybrid samples (Figure S4).

#### Remove bad replicates

We checked the correlation of the read counts between each of the 40 allele-specific expression profiles (function cor() in R (R Foundation for Statistical Computing, Vienna, Austria., n.d.), Figure S5), and found that the expression profiles from two replicates 14A and 9A are apparently different from other replicates. These two replicates are happened to be the two outliers in Figure S4 (the leftmost and rightmost point). We decided to remove them for downstream analysis.

### *Cis* variation estimation

We use the ratio of two alleles’ expression in the hybrid to measure cis variation e_cis_ between the two alleles in one gene.

#### Binomial model

If one assumes that the read counts in a gene for two alleles X and Y in one sample can be modeled by independent Poisson Variables, X and Y can be expressed as:

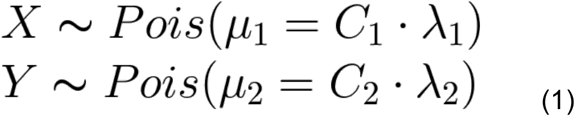

C1 represents the total read counts from one genotype which X allele rested on. C2 represents the total read counts from the other genotype which Y allele rested on. In true hybrid samples, C1 and C2 are almost the same, but in simulations or parental samples they’re not necessarily the same; λ_1_ and λ_2_ represent the proportion of reads mapping to the corresponding alleles. The total read counts C1 and C2 are variable across biological replicates, while λ_1_,λ_2_ are assumed to be biological properties of a gene (expression level) that keep constant across biological replicates. The *cis* effect (e_cis_) is related to the mapping rate parameter λ as follows:

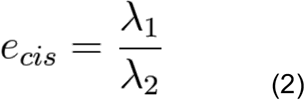

Conditionally on X+Y = n, the probability of k reads mapped to X allele (X=k) is:

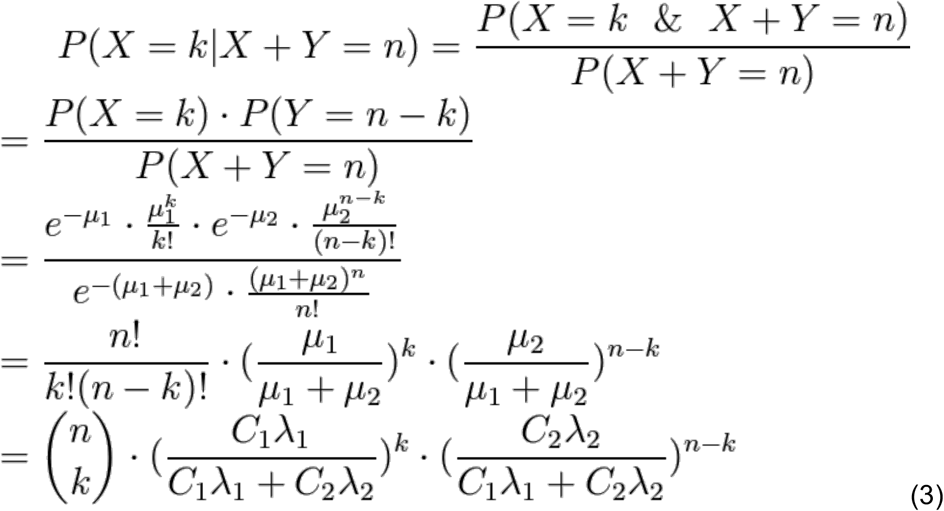

So, the read counts of X allele can be modeled by a binomial distribution conditioned on the sum of the two alleles:

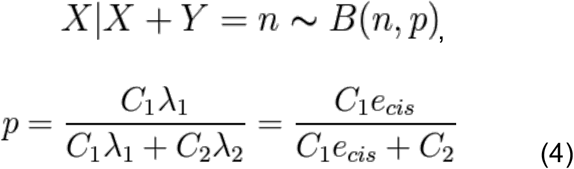

The pdf (probability density function) for X allele’s count in one sample is

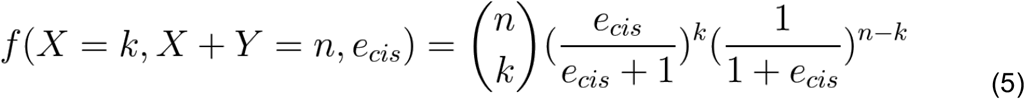

Since the reads count variable X is independent across t biological replicates, the joint pdf is the product of the above pdf. Thus, we can use the Maximum likelihood method to estimate e_cis_. The log-likelihood function to maximize is

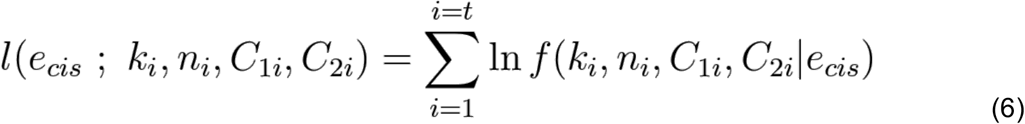

For accommodating the mel2() function in R, in which we applied the log-likelihood function, the optimization for e_cis_ is done on log space. The output is log_2_(e_cis_) and its confidence interval.

#### Beta-binomial model

The assumption that the read counts for alleles can be modeled by independent Poisson Variables may not be appropriate since there is usually more variability than the Poisson Model.

The negative-binomial model provides a good fit to the gene-level read counts distribution (Robinson and Smyth 2007). It’s equivalent to the gamma-Poisson model where the Poisson rate is gamma distributed, adding one degree of freedom to adjust the variance independently of the mean. We now use negative-binomial variables to model the read counts mapped to allele X and Y in the hybrid sample.

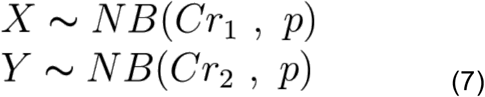

The mean, variance and variance-to-mean ratio for X and Y are shown below:

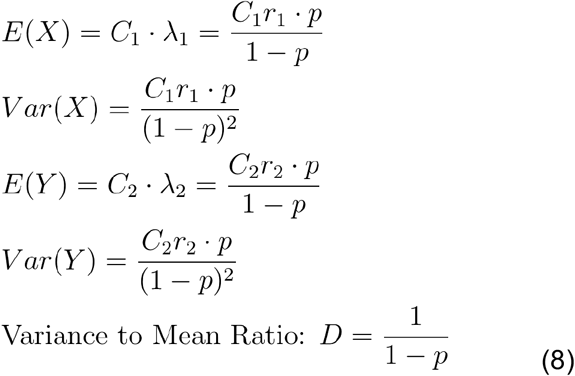

C_1_ and C_2_ represent the total read counts for each genotype in the sample as in binomial model; λ_1_ and λ_2_ represent the proportion of reads mapping to the corresponding alleles; λ_1_=r_1_*p/(1-p) and λ_2_=r_2_*p/(1-p). The assumption for the above modeling is that the two alleles of the same gene have the same variance-to-mean ratio D (p is a constant for X and Y). It is necessary for deriving the beta-binomial distribution below. Although this assumption can not reflect reality completely, it’s still more relaxed than the previously used Poisson model in which the variance equals the mean. When p approaches 0, the negative-binomial model approaches the Poisson Model.

The cis variation e_cis_ is related to the parameter r in the above model:

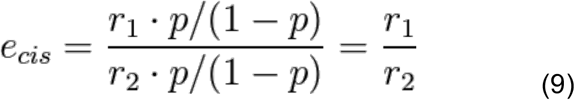

Conditionally on X+Y = n, the probability of k reads mapped to X allele (X=k) is:

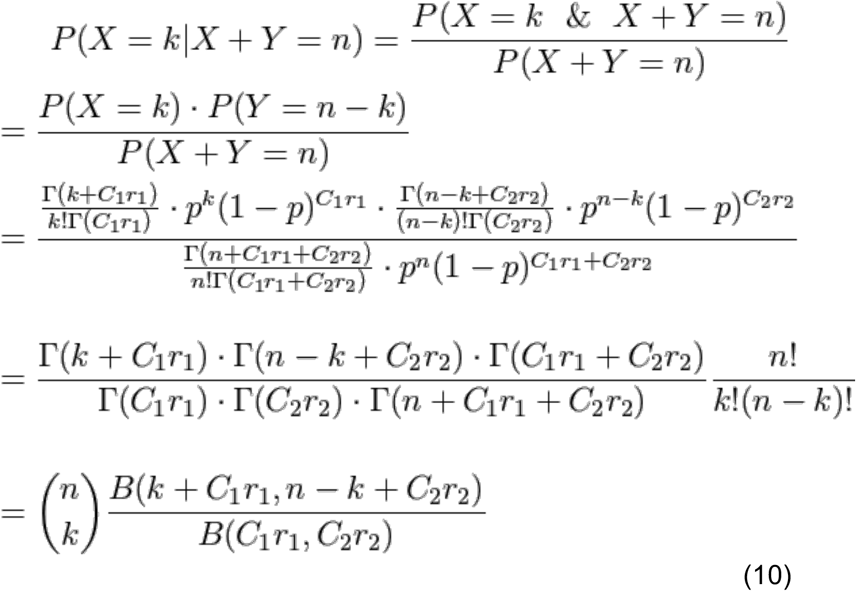

So, the read counts of X allele can be modeled by a beta-binomial distribution conditioned on the sum of the two alleles:

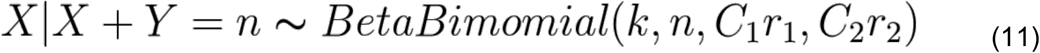

In order to incorporate e_cis_ into the distribution, we reparametrize the beta-binomial distribution with e_cis_ and which describes the over-dispersion of the beta-binomial distribution from the corresponding binomial distribution. Let:

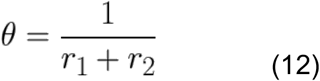

Then from equation 8:

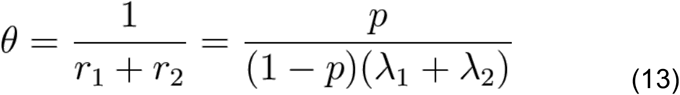

It shows that θ is positively correlated with p.

Then, together with equation(9), we got:

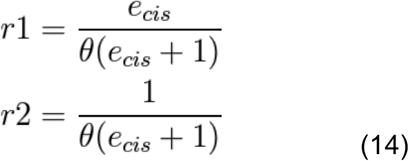

The beta-binomial model approaches the binomial model when θ approaches zero. With the new parameterization, the pdf (probability density function) for X allele’s count in one sample is

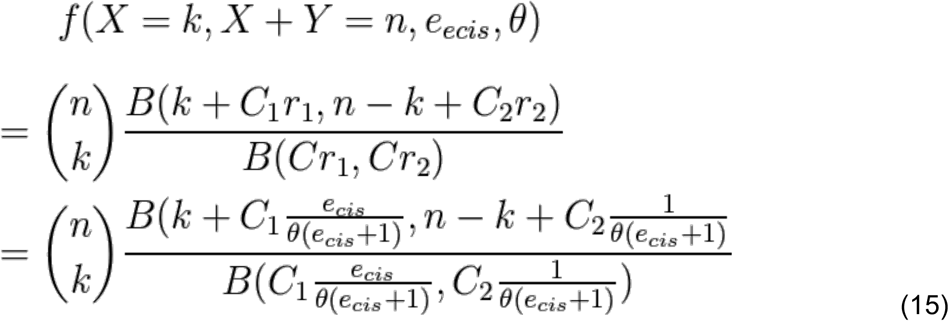

Since the reads count variable X is independent across t biological replicates, the joint pdf is the product of the above pdf. Thus, we can use the Maximum likelihood method to estimate the cis variation e_cis_ along with the over-dispersion parameter θ. The final log-likelihood function to maximize is

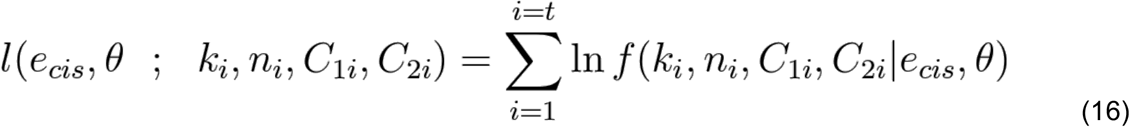

As in the binomial model, the final estimation of e_cis_, θ and their confidence intervals are on log space. The outputs are log_2_(e_cis_) and log_2_(θ) and their confidence intervals.

#### C1 and C2 parameter estimation

For calculating the e_cis_ for a gene, the maximum likelihood method for both models need 4 input from each replicates: k_i_, n_i_, C_1i_, C_2i_.

C_1i_ and C_2i_ are the total expression read counts of the two genotypes. Since there are around 80% of reads in hybrid samples cannot be identified of which genome they belong to, the total allelic reads number cannot be known accurately.

Here we just used the total identifiable read counts from YPS128 allele as C_1_ and those from RM11-1a allele as C_2_ for each sample. That is to say that the aforesaid λ_1_ and λ_2_ are no longer the mapping rate relative to total allelic read counts but to total identifiable allelic read counts. This does not affect the estimation of e_cis_ and its confidence interval If we assume that the identifiable read counts are proportional to true read counts of each allele in the hybrid samples.

### Generate null datasets lacking *cis*-variation

In order to compare the binomial model and the beta-binomial model. We generated two datasets from experimental data which in principle should have no *cis*-variation and four datasets from negative-binomial (gamma-Poisson) distributed random number.

#### Null datasets from experiments

The first dataset “Gier2015” was generated from a haploid yeast gene expression study (Gierliński et al. 2015) which has 48 biological replicates under the same condition: snf2. We downloaded the short reads data from ENA (ENA archive, Project ID: PRJEB5348), then, as described in the paper, got rid of four bad replicates (rep6, rep13, rep25, rep35) and obtained gene read counts with TopHAT2 (Kim et al. 2013) and HTseq (Anders, Pyl, and Huber 2015). We then combined every two haploid expression profiles into 1892 (P(44,2) = 44 × 43) hybrid samples (Table 1).

**Table 1:**
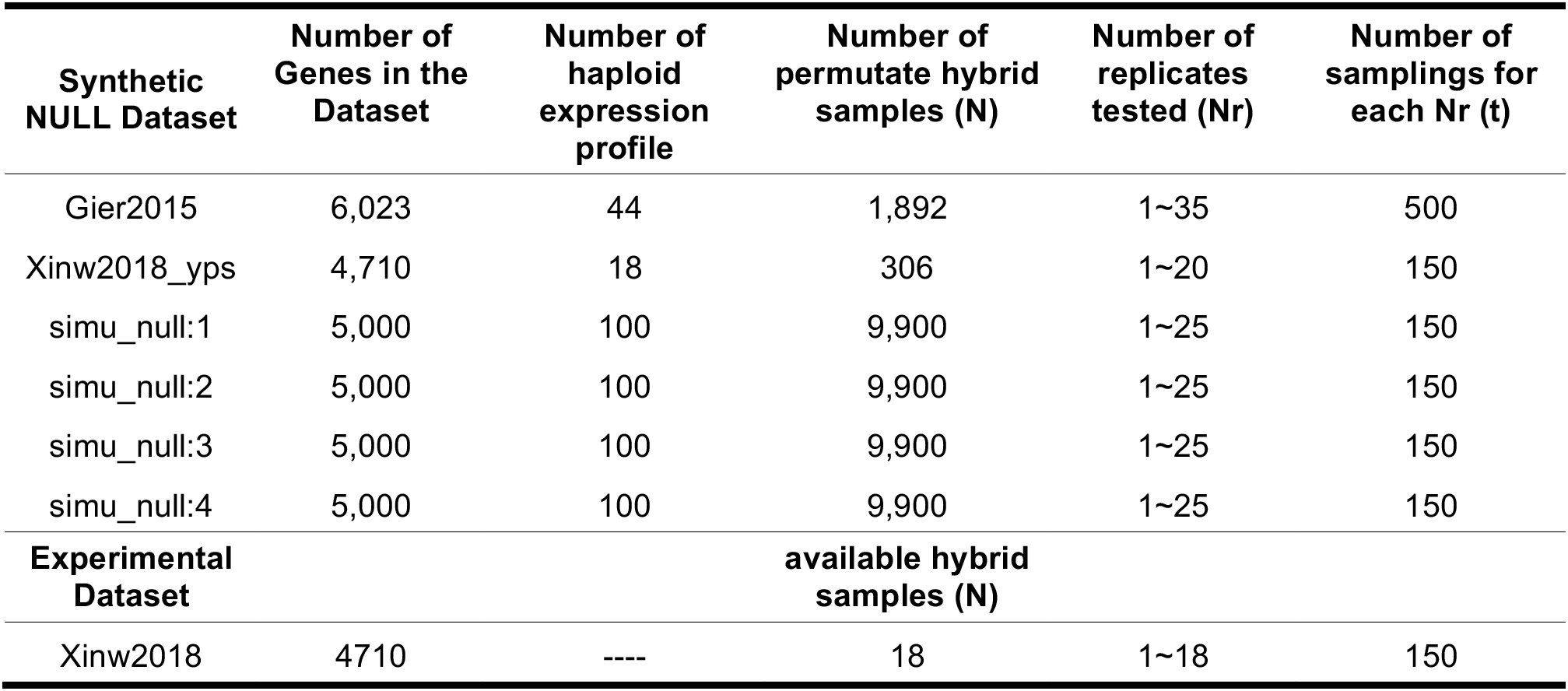
Summary of the datasets used for this study (for details see Materials and Methods: Generate NULL datasets of no cis-variation). Gier2015 and Xinwe2018_yps are null datasets simulated from replicate expression profiles. Simu_null:1-4 are null datasets simulated from random number generator. Xinw2018 are real experimental data.

The second dataset “Xinw2018_yps” was generated in a similar way but from our hybrid samples. We combine every two gene expression profiles from 18 qualified replicates of YPS128 allele, which generated 306 (P(18,2) = 18 *17) no-*cis* hybrid samples (Table 1).

Some simulated hybrid samples have less variation between two alleles, some have more, but by doing this permutation and the bootstrap (see below), the structural bias from choosing extreme hybrids by chance can be attenuated and the average effect of models can be obtained.

#### Null datasets from random number

Although the null dataset generated from experimental data should in principle have no cis-variation, the variation between alleles is not controlled and the true underlying distribution is unknown. So to test both binomial and beta-binomial model with fully-defined hybrid samples, we generated four datasets “simu_null: 1-4” for 5000 genes from the negative-binomial (gamma-Poisson) distribution (Table 1).

The expression counts of each allele for gene i was generated from the negative-binomial distribution using R:

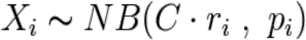

We set C = 1e6; pi was set to 0.1, 0.4, 0.8 respectively for “simu_null:1-3”. For “simu_null:4”, we used a variable p for each gene, which was chosen randomly from a uniform distribution of (0,0.8). We made the gene i have the same expected mapping probability λ across the four datasets, which was chosen by randomly picking a gene from our experimental expression data and use its averaged mapping probability. Since the mapping probability for each gene is set, r_i_ for each gene was then calculated (r_i_ =λ *(1-p)/p) and used as a parameter to generate X_i_.

As a result, each gene across these four datasets have the same expression level, while the variance is getting larger as p_i_ getting larger. Since every two expression profiles were combined to make hybrids within each dataset, there would be no true *cis*-variation. The variance between alleles or among hybrid samples would be low in “simu_null:1” and high in “simu_null:3”.

### Bootstrap cis variation estimation

To test the discovery rate or the false positive rate with different replication number, we randomly choose (without replacement) Nr replicates from all N hybrids. For each level of replication (i.e., Nr), we did the resampling from these N hybrids for t times. Each time, we calculated e_cis_ and its 95% confidence interval using maximum likelihood method (See Method: Cis variation estimation). If a gene’s log_2_(e_cis_) confidence interval overlap with 0 (e_cis_ = 1), we classify it as a significant cis-variant gene. Table 1 shows the N (number of hybrid samples), Nr (Number of replicates tested), and t (number of samplings) for each dataset.

## Results

### Assembly of reference genomes

Read mapping biases related to using only a single reference genome will lead to biases in allelespecific expression inference (Degner et al. 2009). To mitigate such bias, we constructed two reference quality *de novo* genome assemblies of the parental strains used in this study, YPS128, and RM11-1a.

The contiguity, completeness, and accuracy of our assemblies are quite high (Table 2 and Figure 1). Both assemblies exhibit a high level of contiguity, with the majority of chromosomes being covered by one or two contigs, comparable to that of the *Saccharomyces cerevisiae* S288C Reference R64-1-1 (Table 2 and Figure 1). The BUSCO score assesses genome assembly completeness by identifying conserved single copy orthologs (Simão et al. 2015). Both assemblies compare favorably to the yeast community reference genome (Table 2: RM11-1a: 93.7%; YPS128: 94.4%; R64: 93.9%). The QV scores we calculate reflect the basepair-level concordance between an assembly and Illumina short reads (Koren et al. 2018). While the new assemblies are both quite accurate, due to the lower coverage and noisier long reads used in assembling RM11-1a, its assembly exhibited a lower QV even after polishing (Table 2: RM11-1a: 35.6; YPS128: 60.0).

**Figure 1:**
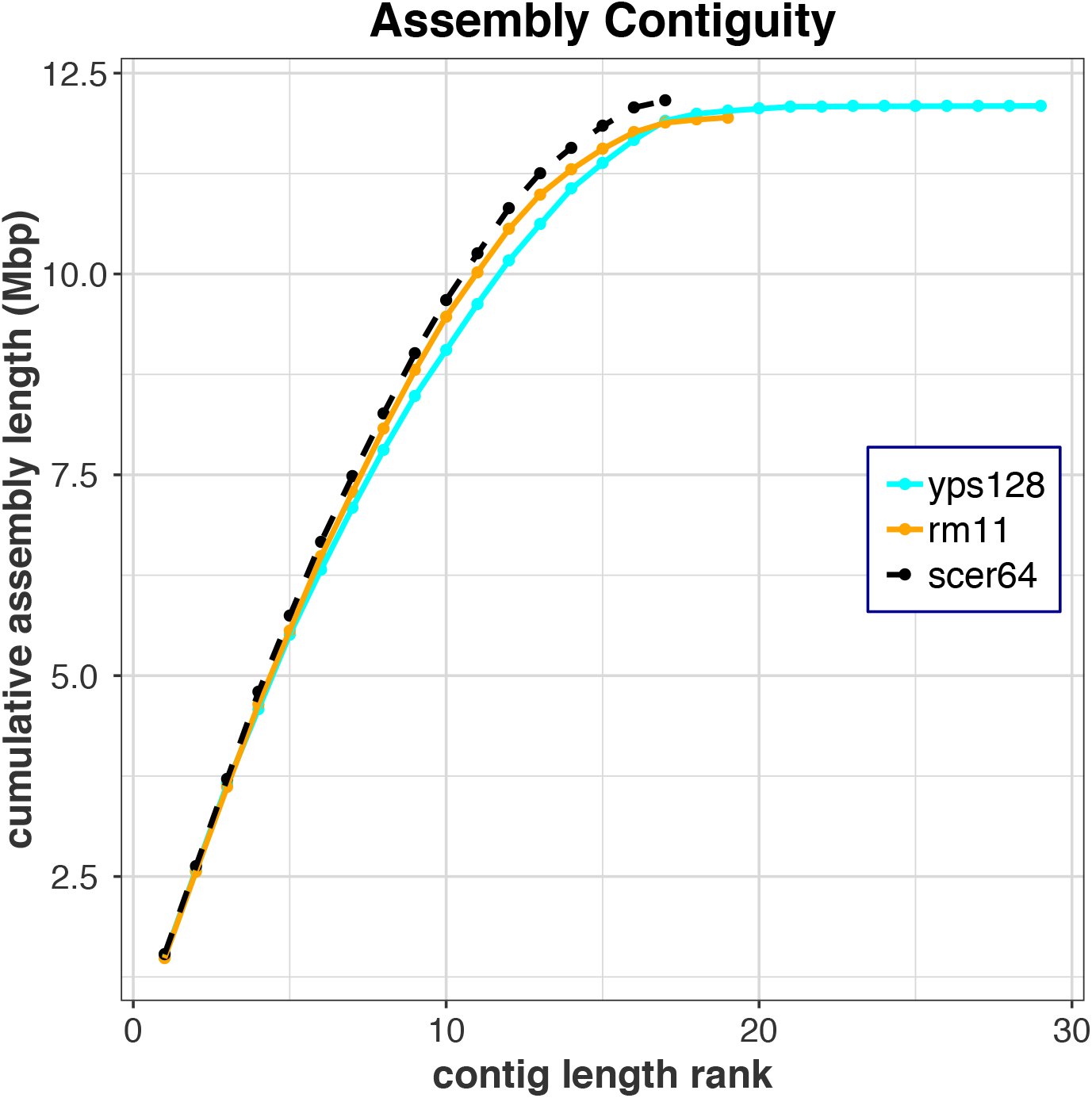
The contiguity of our assemblies is comparable to that of the Saccharomyces cerevisiae S288C Reference R64-1-1. Contigs are ranked from longest to shortest. Their cumulative sum of length are shown on Y axis in mega bases.

**Table 2:** The contiguity, completeness, and accuracy of YPS128 and RM11-1a genomes.

|  | S.cer R64-1-1 | Rm11-1a | Yps128 |
| --- | --- | --- | --- |
| Assembly size (Mb) | 12.16 | 11.95 | 12.09 |
| number of Contigs/scaffolds | 17 | 19 | 29 |
| Contig N50 (Mb) | 0.92 | 0.92 | 0.81 |
| Contig L50 | 6 | 6 | 6 |
| Contig N90 (Mb) | 0.44 | 0.43 | 0.44 |
| Contig L90 | 13 | 13 | 14 |
| Busco score | 93.9% | 93.7% | 94.4% |
| Complete Busco | 1,351 | 1,347 | 1,358 |
| Fragmented Busco | 38 | 37 | 33 |
| Missing Busco | 49 | 54 | 47 |
| QV score | ---- | 35.6 | 60.0 |

### Allele-specific RNAseq

We sequenced 20 replicates of hybrid mRNA samples. The 20 samples were used independently to construct 20 barcoded libraries that were pooled into a single sequencing experiment. After demultiplexing, we obtained 1.531 billion 75-bp paired-end reads. We then counted the allelespecific counts for each gene using the SNPs/Indels between YPS128 and RM11-1a genomes. Mapping bias was eliminated by using both YPS128 and RM11-1a genomes as references in the mapping step and filtering out suspect SNPs/indels (Figure S4). We then discarded two replicates exhibiting the lowest correlation with other replicates (Figure S5), and finally obtained 18 replicates of allele-specific gene read counts for 4,710 genes.

### The beta-binomial distribution models cis-expression sampling variation better than the binomial distribution

To assess the performance of two models, we additionally simulated 6 hybrid null datasets lacking true *cis*-variation (for details, see Materials and Methods: Generate null datasets lacking *cis*variation & Table 1). For each dataset (Table 1), we applied our inference machinery to estimate the *cis*-variation parameter and its 95% confidence interval for each gene. As the null data exhibits no true *cis*-variation, any significant expression should be caused by false positives. We then plot the rate of rejecting null hypothesis (which reduced to the false positive rate in the null simulations) against replication to examine the behavior of the models as power increases (Figure 2–4). The beta-binomial model exhibited a false positive rate closer to the prediction than the binomial model in null datasets. However, for both models, the performance was poor when there was little replication (Figure 2–3).

**Figure 2:**
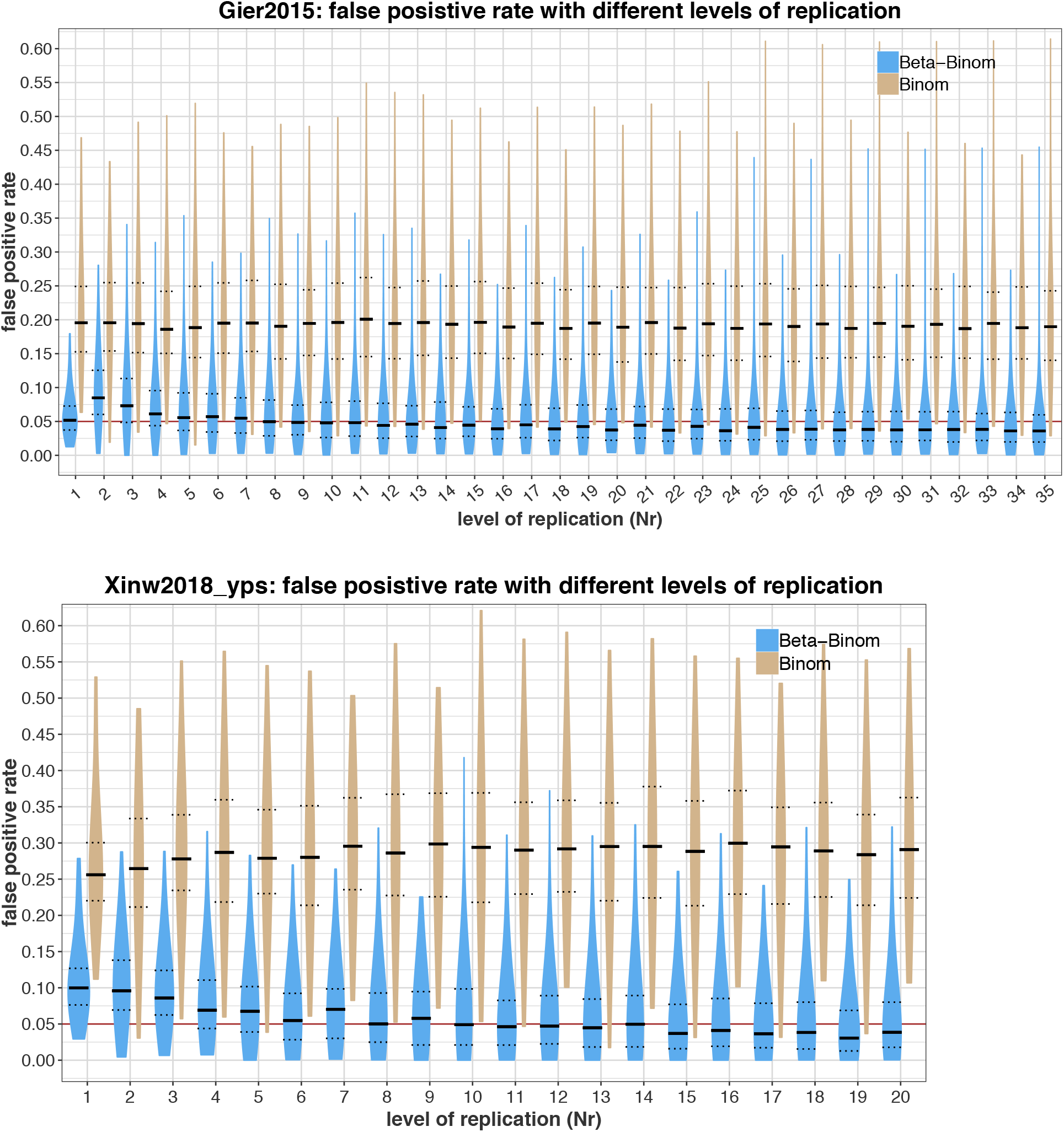
False positive rate with different number of replicates. The blue and brown violin plot in each level of replication show the distribution of false positive rates from t (See Table 1) sampling results. The red horizontal line is the expected false positive rate of 0.05 (α = 0.05). The solid and dot lines on each plot are the median, 25% quantile and 75% quantile. A) The binomial model consistently rejects the null hypothesis at a rate around 20%. The betabinomial model consistently exhibits a rejection rate that is lower than that of the binomial model and is getting closer to the expected 5% as more replicates used. The rejection rates of betabinomial model improve with sufficient replication, approaching the significant level α. B) The binomial model consistently rejects the null hypothesis at a rate around 28%. Similar to panel A, the beta-binomial model consistently exhibits a lower rejection rate and is getting closer to the expected 5% as more replicates used.

**Figure 3:**
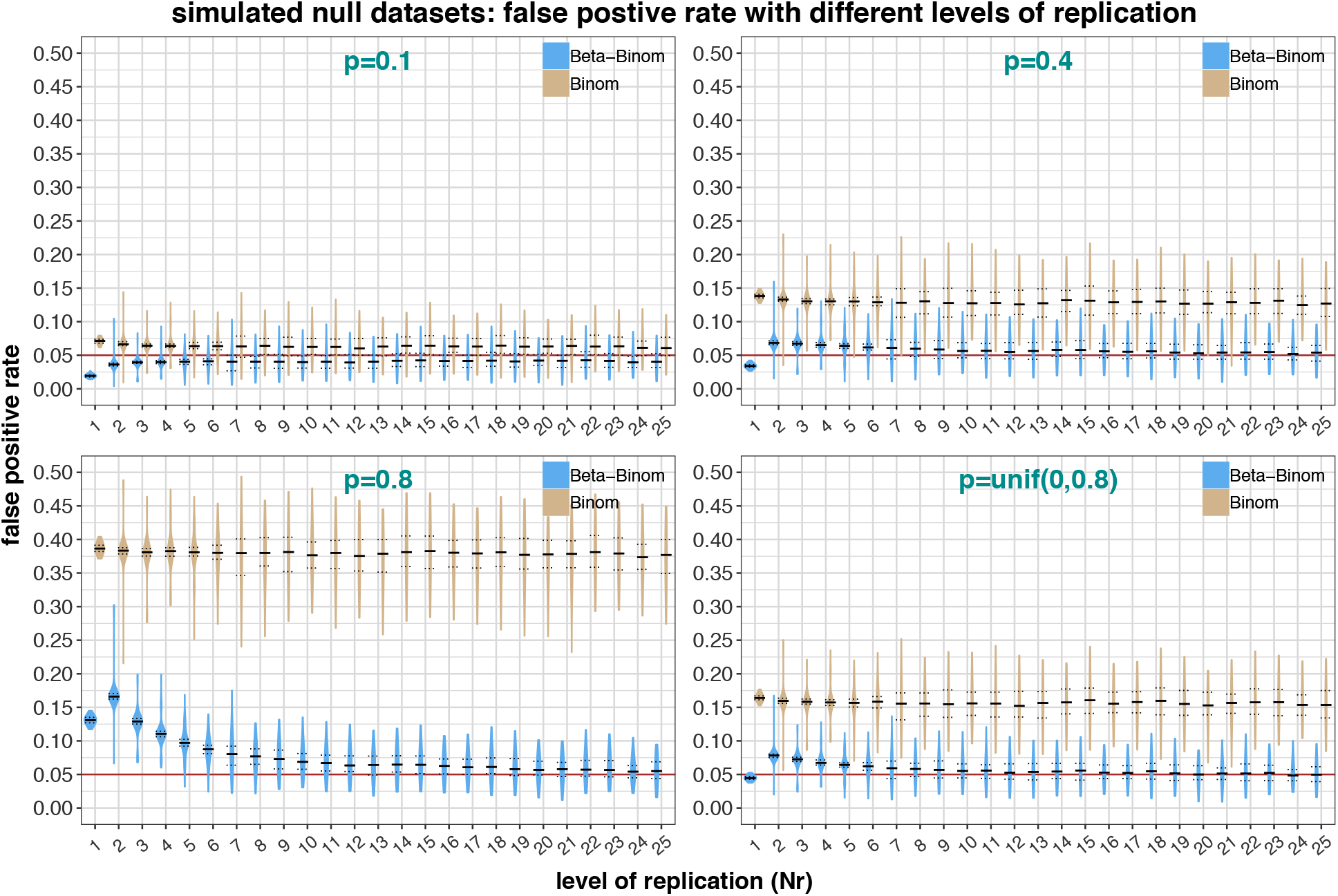
False positive rate with different number of replicates in simu_null:1-4. The degree of excess false positives is related to the simulated over-dispersion of each dataset (p is a parameter controls the over-dispersion: when p approaches zero, the gamma-Poisson model approaches the Poisson model; when p approaches one, the gamma-Poisson model is strongly overdispersed). The binomial model has a consistent ~7% false-positive rate in “simu_null:1” which is only 2% higher than expected 5% (panel p=0.1), but it can be as high as 38% in “simu_null:3” (panel p=0.8). The performance of the binomial model on a changing “p” (panel p=unif(0,0.8)) is similar to the constant “p” with the corresponding mean except a ~3% more false-positives (panel p=0.4). With few replicates and low overdispersion, the beta-binomial demonstrates a lower falsepositive rate than expected (panel p=0.1). The reverse is true under the high overdispersion simulation. The beta-binomial model consistently approaches α with increasing replicates.

**Figure 4:**
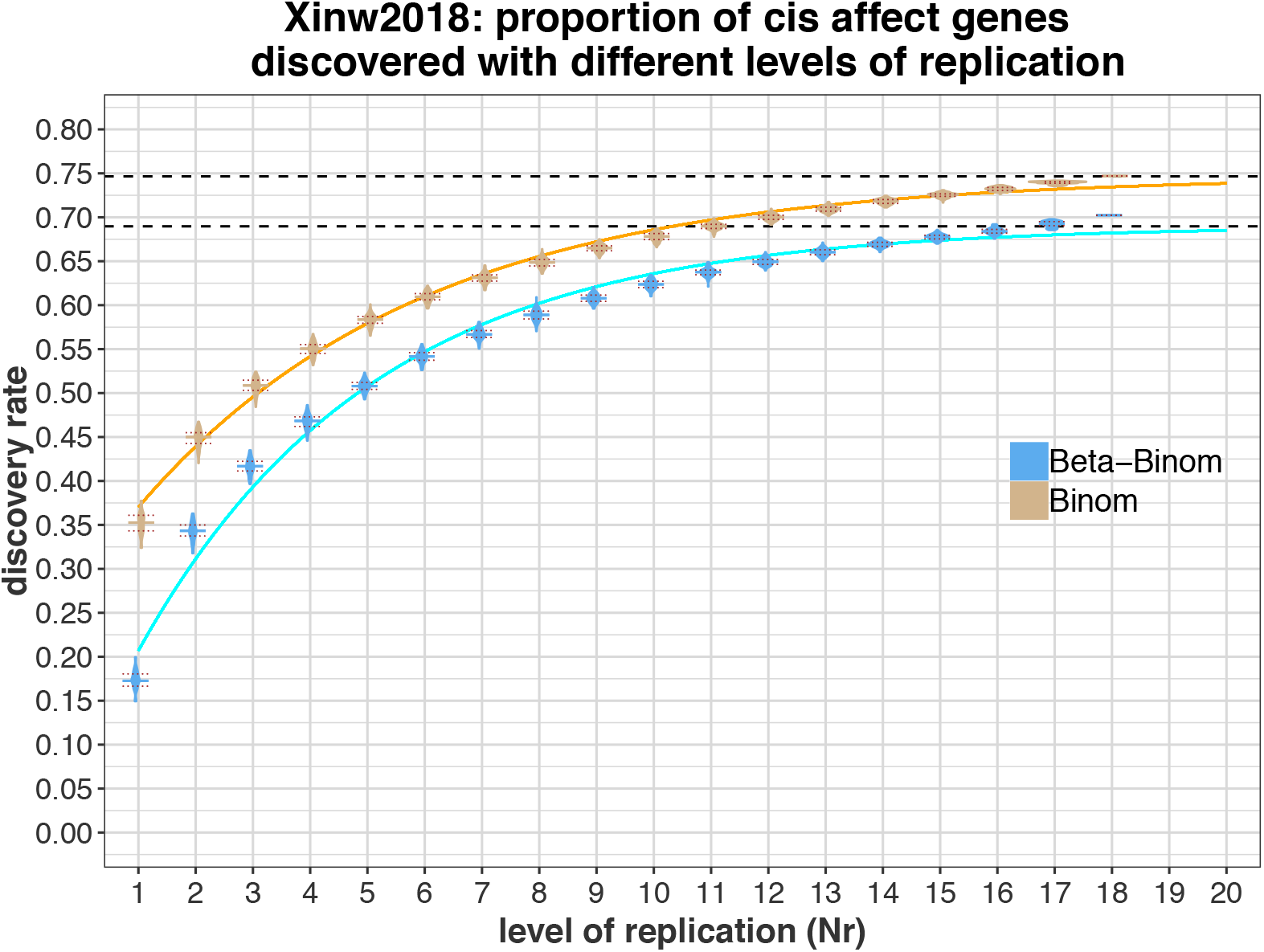
The discovery rate of the dataset “Xinw2018”. More significant genes are discovered as more replicates used. When all 18 replicates are used, the rate of rejection appears to asymptote to ~70% for the beta-binomial model and 75% for the binomial model. The two asymptotic lines were drawn by fitting the data to a negative exponential model.

### Inference on a highly replicated dataset without genetic or environmental variation

One validation of our model makes use of data from a yeast expression experiment comprising 44 biological replicates of a single haploid yeast strain under the same condition (Gierliński et al. 2015). Pairs of expression profiles were combined into synthetic/*in silico* hybrid samples by permuting pair assignments such that the two alleles within one synthetic hybrid do not have any true *cis* variation while retaining the sample variation between them (for details, see Materials and Methods: Generate null datasets lacking *cis*-variation & Table 1). This hybrid dataset was labeled “Gier2015”, yielding 1,892 permuted synthetic hybrid samples (44*43 = 1,892).

To test whether increasing replication improves *cis* estimation, we randomly sampled Nr replicates without replacement from the 1,892 synthetic hybrids, performed *cis* parameter inference, and calculated the false positive rate. Nr ranged from 1 to 35 for this dataset. For each level of replication (i.e., Nr), we sampled, as described above, 150 times to determine the distribution of the false positive rate (Figure 2A; see Materials and Methods: Bootstrap *cis* variation estimation).

We generated and analyzed another hybrid dataset “Xinw2018_yps” following a similar approach to “Gier2015”, but using our own expression experiment. The 18 expression profiles of the YPS128 allele were extracted from the 18 hybrid samples (2 of the 20 replicates are removed due to being outliers as measured in terms of exhibiting low correlation with other replicates), and pairs of profiles were combined into 306 (18*17=306) synthetic hybrids (Figure 2B).

Our results demonstrate that the binomial model consistently rejects the null hypothesis at an elevated rate for α = 0.05, exhibiting a consistent rejection rate across levels of replication (Figure 2A-B). The beta-binomial model consistently exhibits a rejection rate that is lower than that of the binomial model. However, the beta-binomial does show some variation in rejection rate at low replication. In particular, for low replication in both the “Gier2015” and “Xinw2018_yps” datasets, the beta-binomial model shows an excess rate of rejection that subsides as replication increases.

The severity in underestimating variance using the binomial model depends on the underlying variance among replicates. The rejection rate of the binomial model can vary from 20% (Figure 2A) to as high as 30% (Figure 2B). Increased replication seems to have little effect on diminishing this problem. In contrast, the false-positive rate in the beta-binomial model improves as replication increases. The increasing of the false-positive rate at the beginning of Figure 2A is likely an artifact resulting from the starting point of the maximum likelihood optimizer (See Materials and Methods: *Cis* variation estimation). Other than this artifact, the beta-binomial model also appears to underestimate the variation among replicates with fewer replicates, leading to high false-positive rate, though reduced as compared to the binomial model. The rejection rates improve with sufficient replication, asymptoting towards the significant level α.

### De novo null simulation

Although the null datasets we generated by randomly pairing real experimental replicates exhibit no true *cis* variation, there is the potential for unknown confounding factors that were not controlled. We therefore simulated four hybrid datasets for 5,000 genes from the gamma-Poisson distribution (“simu_null: 1-4”), with the same expression level between alleles and explicit overdispersion parameters so that we can study the behavior of overdispersed expression data in the absence of differential gene expression.

The gamma-Poisson distribution (also known as the negative-binomial distribution) is widely used to model the read counts distribution among replicates (Robinson and Smyth 2007, 2008). This distribution can be viewed as a Poisson distribution where the Poisson parameter is gamma distributed.

We simulated the expression profile of 5,000 genes across a wide number of expression levels under this model. The four different datasets with different over-dispersion profiles were generated by systematically varying the “p” parameter in the gamma-Poisson distribution for each dataset. We ensured that each gene maintained the same expression level across all four datasets. When p approaches zero, the Gamma-Poisson model approaches the Poisson model. When p approaches one, the Gamma-Poisson model is strongly over-dispersed (for details, see Materials and Methods: Generate null datasets lacking *cis*-variation: Null datasets from random number). We randomly paired samples within each of the four datasets following the same approach described above for “Gier2015” and “Xinw2018_yps”. This permitted us to vary the level of overdispersion and study the consequences for inference.

We set the p parameter to 0.1, 0.4, and 0.8 for the first three datasets (simu_null:1-3 respectively). As a result, the first dataset (simu_null:1) has the lowest over-dispersion with expression profiles (the closest to the Poisson model) whereas the third (simu_null:3) is the most over-dispersed. For the final simulation (simu_null:4) we chose a uniform distribution of p parameters with a mean of 0.4 for the 5,000 genes to simulate the impact for genome-wide inference when a dataset has genes with different levels of overdispersion.

The false-positive rate for these four datasets shows a similar pattern as in “Gier2015” and “Xinw2018_yps”. The binomial model shows an elevated false-positive rate that is not mitigated with increased replication (Figure 3A-D). The degree of excess false positives is related to the simulated over-dispersion of each dataset. The binomial model has a ~7% false-positive rate in “simu_null:1” which is only 2% higher than the expected 5% (Figure 3A), but it can be as high as 38% in “simu_null:3” (Figure 3C). The performance of the binomial model on a changing “p” (Figure 3D, fp ~16%) is similar to the constant “p” with the corresponding mean with an ~3% higher rate of false-positives (cf. Figure 3B, fp ~13%).

With few replicates and low overdispersion, the beta-binomial demonstrates a lower false-positive rate than expected (Figure 3A), suggesting that it is overestimating the variance when only a few replicates are used. The reverse is true under the high overdispersion simulation, suggesting it is underestimating the variance (Figure 3C). However, the model consistently approaches α with increasing replication. This is likely because the overdispersion parameter θ is poorly estimated with only a few replicates and relies on the initial arbitrary value in maximum likelihood optimizing, a situation that improves with higher replication.

### The effects of replication on ASE confidence intervals

We then applied the e_cis_ inference machinery on the experimental dataset of our 18 replicated hybrid samples (Xinw2018). More significant genes are discovered as more replicates are used. When all 18 replicates are used, we observe the rate of rejection appears to asymptote to ~70% with the beta-binomial model. The number of significant genes from the binomial model exceeds the beta-binomial by ~5% (Figure 4).

To explore the effect of gene expression level and number of replicates on the power, we chose 100 typical genes from each of the following categories: “lowly expressed genes” (average counts: 50-200); “intermediate expressed genes” (average counts: 400-600); and “highly express genes” (average counts: 1500-3500). We plotted the confidence intervals for each gene using the estimation calculated from four levels of replication (3,6,12,18). Genes are ranked by their *cis* effect (Figure 5).

**Figure 5:**
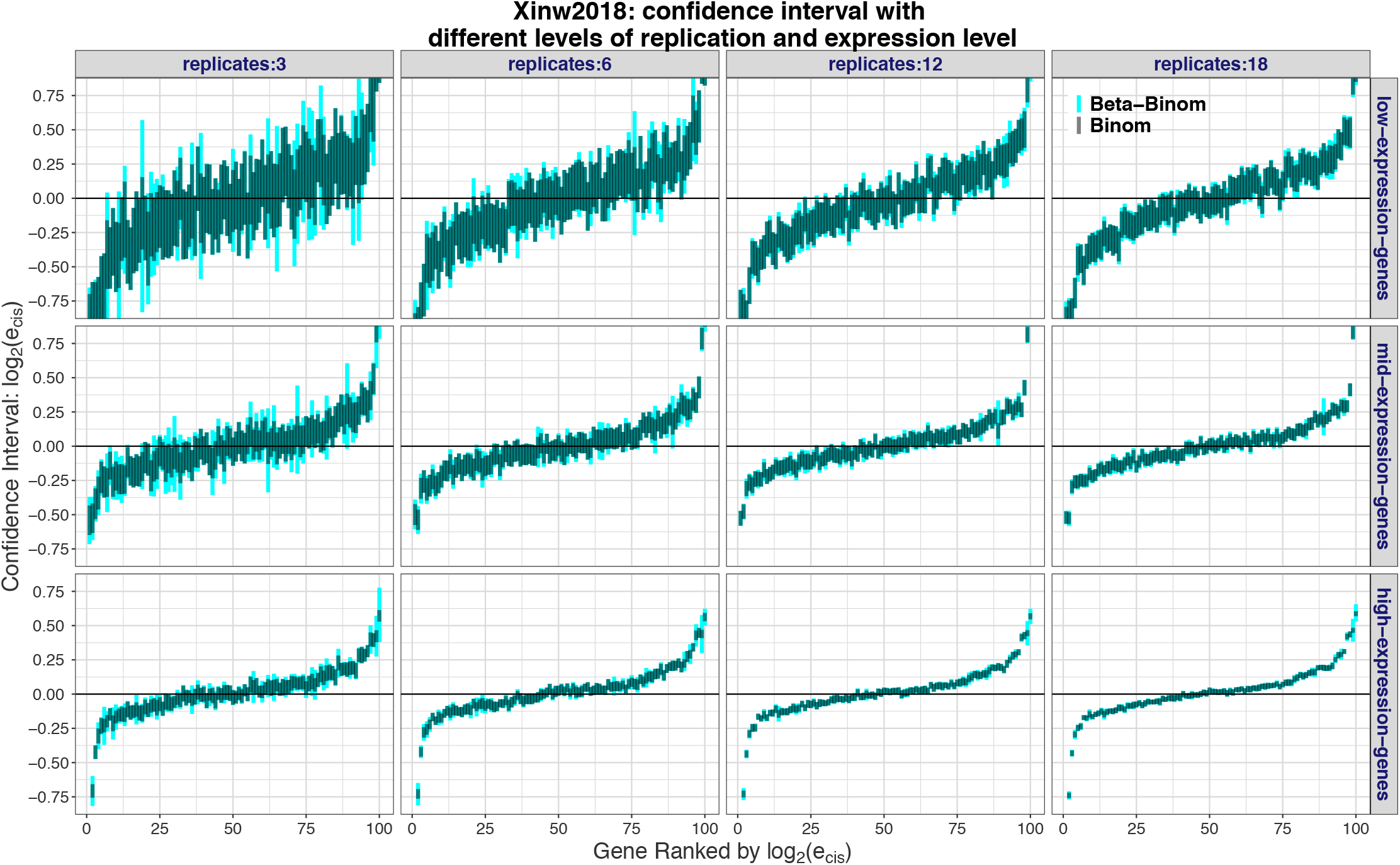
The effect of gene expression level and number of replicates on the inference power. 100 typical genes of low expression (average counts: 50-200), mid expression (average counts: 400-600) and high expression (average counts: 1500-3500) are plotted for their confidence intervals using the estimation calculated from four levels of replication (3,6,12,18). The genes are ranked by their cis-effect: log_2_(e_cis_). Grey is for binomial model; Blue is for beta-binomial model.

As expected, the beta-binomial model yields a wider confidence interval than the binomial model, reducing the false-positive rate. We also see that, as expected, when replication increases or with higher expression level, the confidence intervals narrow for both models, increasing the power (Figure 5).

### *Cis* variation between YPS128 and RM11-1a strain is ubiquitous and often small in magnitude

The rate of rejection appears to asymptote to ~70% (Figure 4) with beta-binomial model in the “Xinw2018” dataset, suggesting that ~70% of the 4,710 genes we studied show evidence for expression variation, a marked increase compared to previous observations (Emerson et al. 2010; Schaefke et al. 2013; Metzger, Wittkopp, and Coolon 2017; Tirosh et al. 2009; Artieri and Fraser 2014).

We then summarized the e_cis_ distribution calculated form all 18 hybrid replicates with the betabinomial model (Figure 6A, Table 3). The symmetry of the distribution of log_2_(e_cis_) indicates that there are similar amount of genes affected by cis-regulatory variation in both directions. Approximately 70% of the genes (3,308 out of 4,710) exhibit significant *cis* variation (|log_2_(e_cis_)|> 0; p < 0.05). Notably, the *cis* effect in most of these significant genes is small in magnitude. Of the differentially expressed genes (Figure 6B), 70% exhibit cis variation in the range 0 < |log_2_(e_cis_)| < 0.2, or less than a 1.15-fold difference. The genes with the *cis* variation |log_2_(e_cis_)| > 1 (i.e. a 2 fold difference) only comprise 3% of all significant genes.

**Figure 6:**
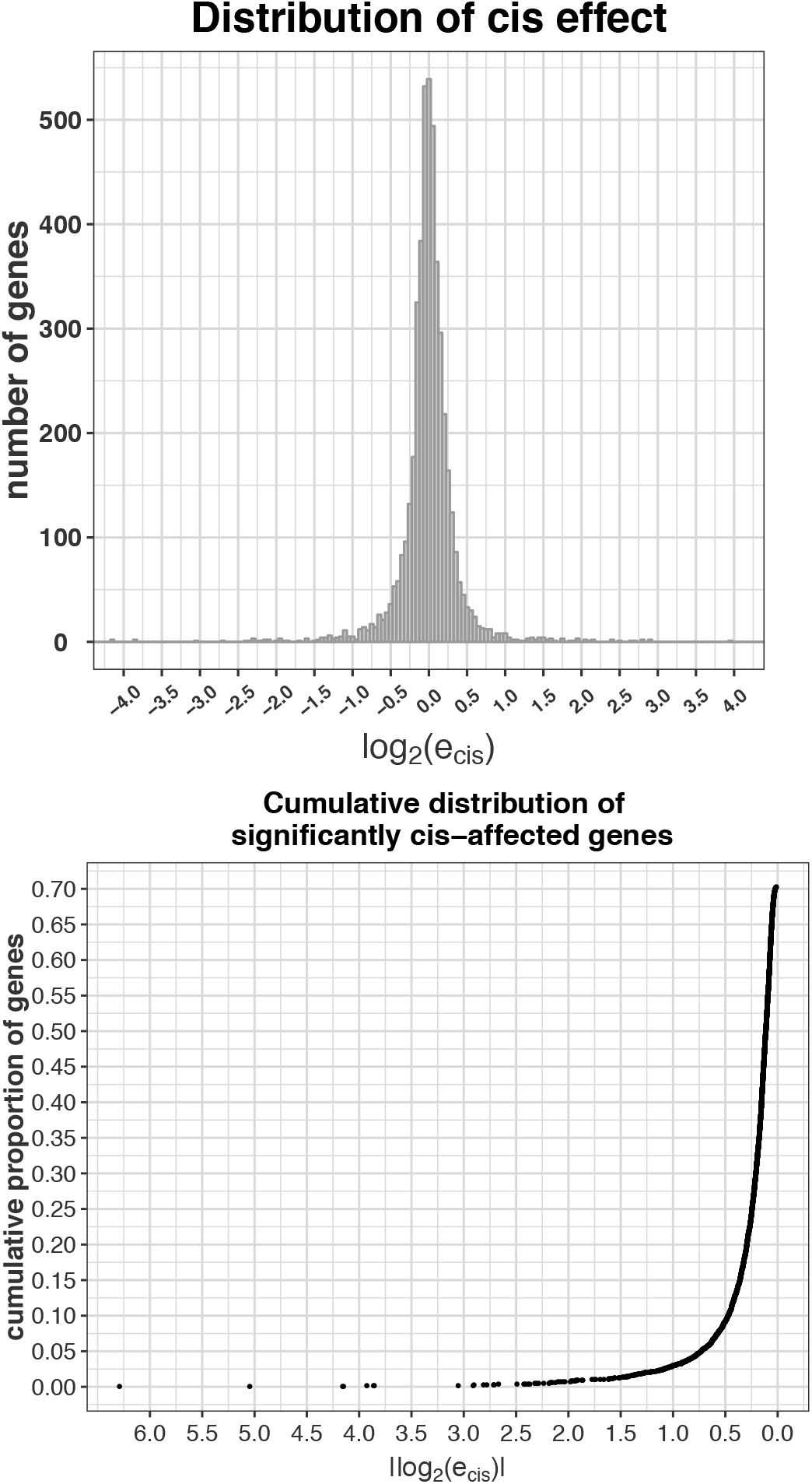
A) The log_2_(e_cis_,) distribution from all 18 hybrid replicates with beta-binomial model. B) The cumulative proportion of significantly *cis*-affected genes. The cis-affected genes are sorted by their cis effect, from largest to smallest. The cumulative proportion shows that most significant genes have a *cis*-effect of small magnitude.

**Table 3:**
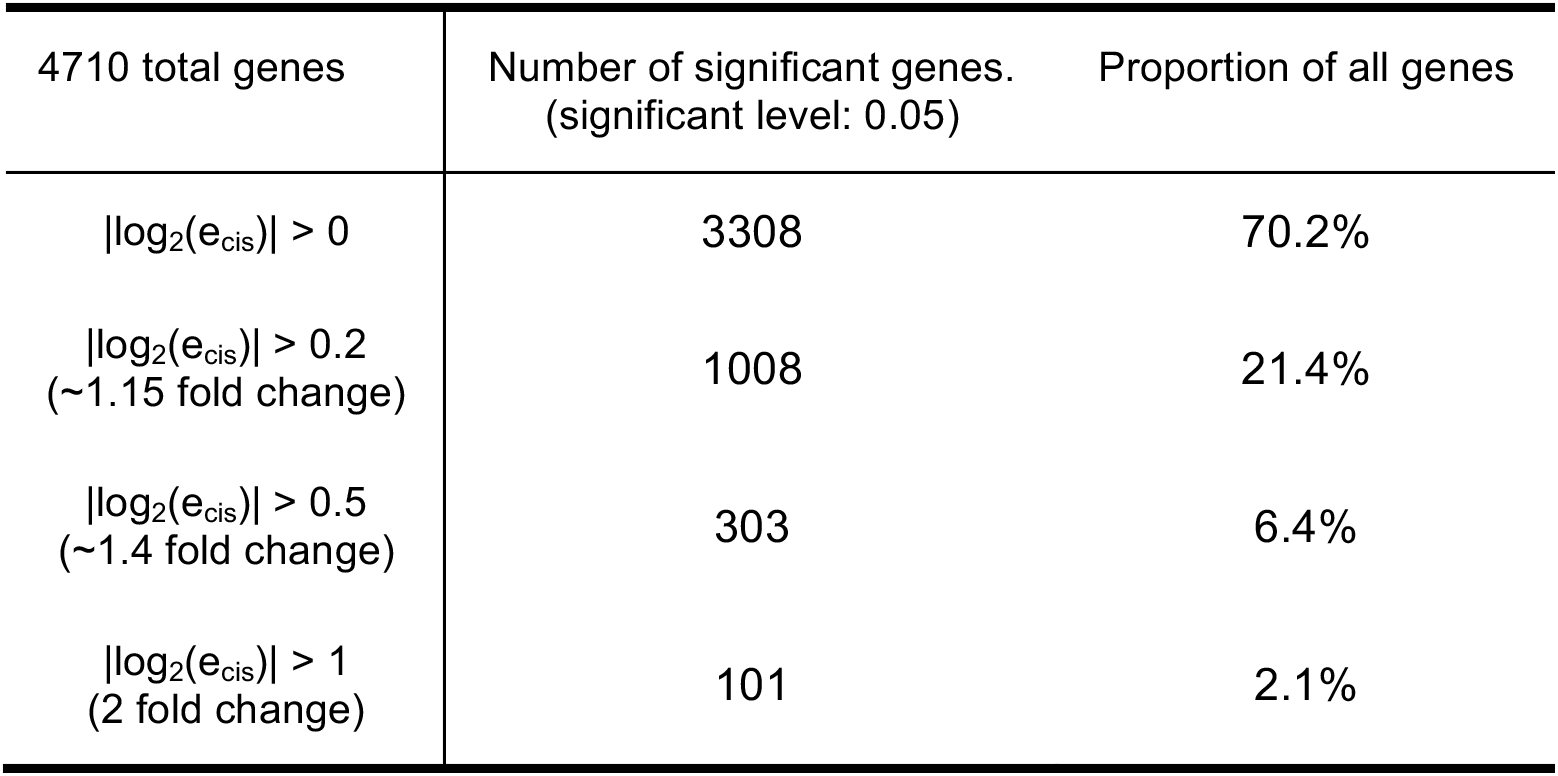
The number of genes and their proportion in different cis-effect magnitude.

## Discussion

Inference of allele-specific expression differences from F1 hybrids is a widely used perspective to explore the evolution of gene expression. Many results have been reported for a wide range of individuals, populations, or species (Tirosh et al. 2009; Wittkopp, Haerum, and Clark 2008; McManus et al. 2010; Emerson et al. 2010). Such inferences have been applied to questions about compensation between *cis* and *trans* variation (Romero, Ruvinsky, and Gilad 2012; Mack, Campbell, and Nachman 2016), stabilizing selection for expression level (Hodgins-Davis, Rice, and Townsend 2015), and *cis*-effect in inter-specific/intra-specific expression variation (Metzger, Wittkopp, and Coolon 2017; Rhoné et al. 2017) and all depend in a central way on accurate measurement of *cis* variation. However, naive statistical models (Robinson and Smyth 2007) and the tendency to misuse replication has limited the utility of allele-specific-expression inference.

In this work, we describe a beta-binomial model for estimation of *cis* expression variation in allelespecific studies. It is based on a more suitable gamma-Poisson distribution of read counts among replicated experiments and is capable of accommodating over-dispersion of expression. We demonstrate the advantage of the beta-binomial model over the binomial model with both experimental and simulated data. The results showed that, with sufficient replication, the betabinomial model attains the nominal false positive rate while the binomial model consistently underestimates the variance leading to an elevated false-positive rate.

While, unlike the Poisson model, the gamma Poisson model permits the variance and mean to be independent, rigorous inference using the beta-binomial model derived from it still requires each allele to exhibit approximately the same variance-to-mean ratio (See Materials and Methods: *Cis* variation estimation: Beta-binomial Model). This limitation can be addressed by assigning different over-dispersion parameters for each allele, but inference becomes more complex. In any event, the good performance of the beta-binomial model suggests that potential improvement for e_cis_ estimation is limited.

The trade-off between the false-positive rate and power still holds in these two models. We used the significant gene list from our best estimates (i.e. the beta-binomial model with all 18 replicates) as a gold standard to explore the relative power of both models (Figure S6 A). The binomial model has higher power than the beta-binomial model in all levels of replication. Of course, even the best statistical model would by definition exhibit α×100% false positives. If we assume the 18 replicate beta-binomial model has 100% of power (Figure S6), then the proportion of true negatives that yields a false positive rate of 0.05 is (1-0.702)/(1-0.05) = 0.314. The 18 replicate binomial model rejects the null hypothesis 74.5% of the time, implying its false positive rate is 18%(0.18=1-(1-0.745)/0.314, assuming 100% power), which is consistent with our simulations (Figure 2A).

We uncovered many more *cis*-affected genes than previous intra-specific studies of yeast, where the proportion varies between 6%-29% (Emerson et al. 2010; Schaefke et al. 2013; Metzger, Wittkopp, and Coolon 2017). The main culprit is likely lower power in previous studies, although we also used YPS128 rather than the BY4741 strain common in previous studies. Figure 4,5 & S6 demonstrate that adding more replicates increases the power and the relative difference in the discovery rate can be as high as 55% (Figure 4). Results from previous studies using one or two replicates yield comparable numbers of genes differentially expressed in *cis* (Figure 4, the left-most two points of the Binomial model), suggesting that the difference in our results is of higher power to detect smaller magnitude changes.

Our results quantify the advantage of the beta-binomial model over the binomial model in detecting *cis* variation. The beta-binomial model estimates variance accurately and also has high statistical power as long as sufficient replicates are provided. Thus, our high replicate experiment describes an accurate and complete landscape of *cis* variation between YPS128 and RM11-1a. We recommend a beta-binomial model should for use in future allele-specific experiments and predict it will reveal an abundance of *cis* variation that previously remained hidden.

The number of genes and number of replicated expression profiles (except Xinw2018, hybrid samples do not have haploid expression profile) are in column 2 & 3. The permuted hybrid samples are listed in column 4. The number of replicates are listed in column 5. The ranges were chosen somewhat arbitrarily, but were enough to see the trend. For level of replication, we did the resampling t times shown in column 6.

## Supplementary Figures

**Figure S1:**
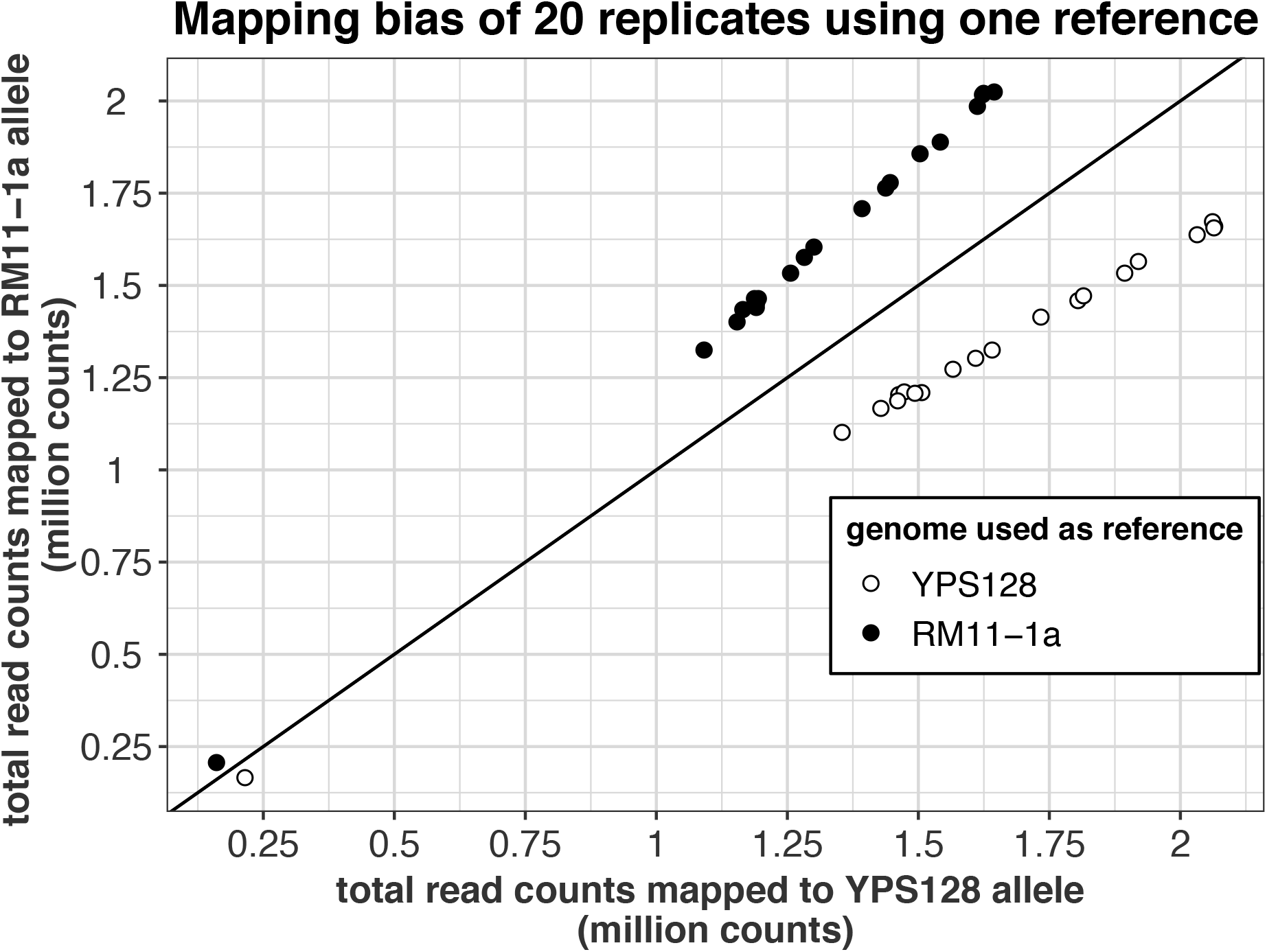
The mapping bias of DNA read counts when using only one assembly as the reference genome.

**Figure S2:**
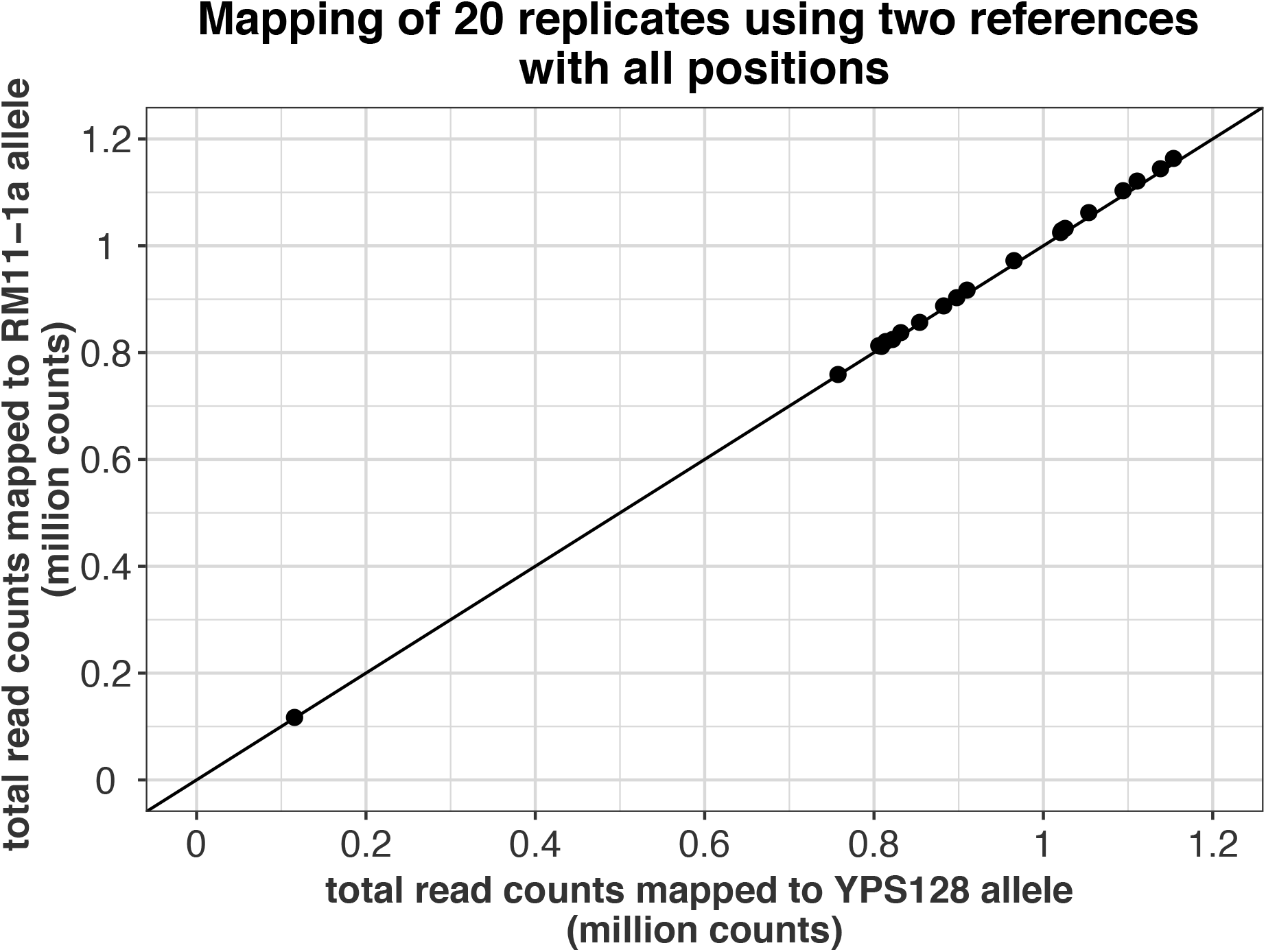
The mapping bias are mostly removed when using both assemblies as reference genomes.

**Figure S3:**
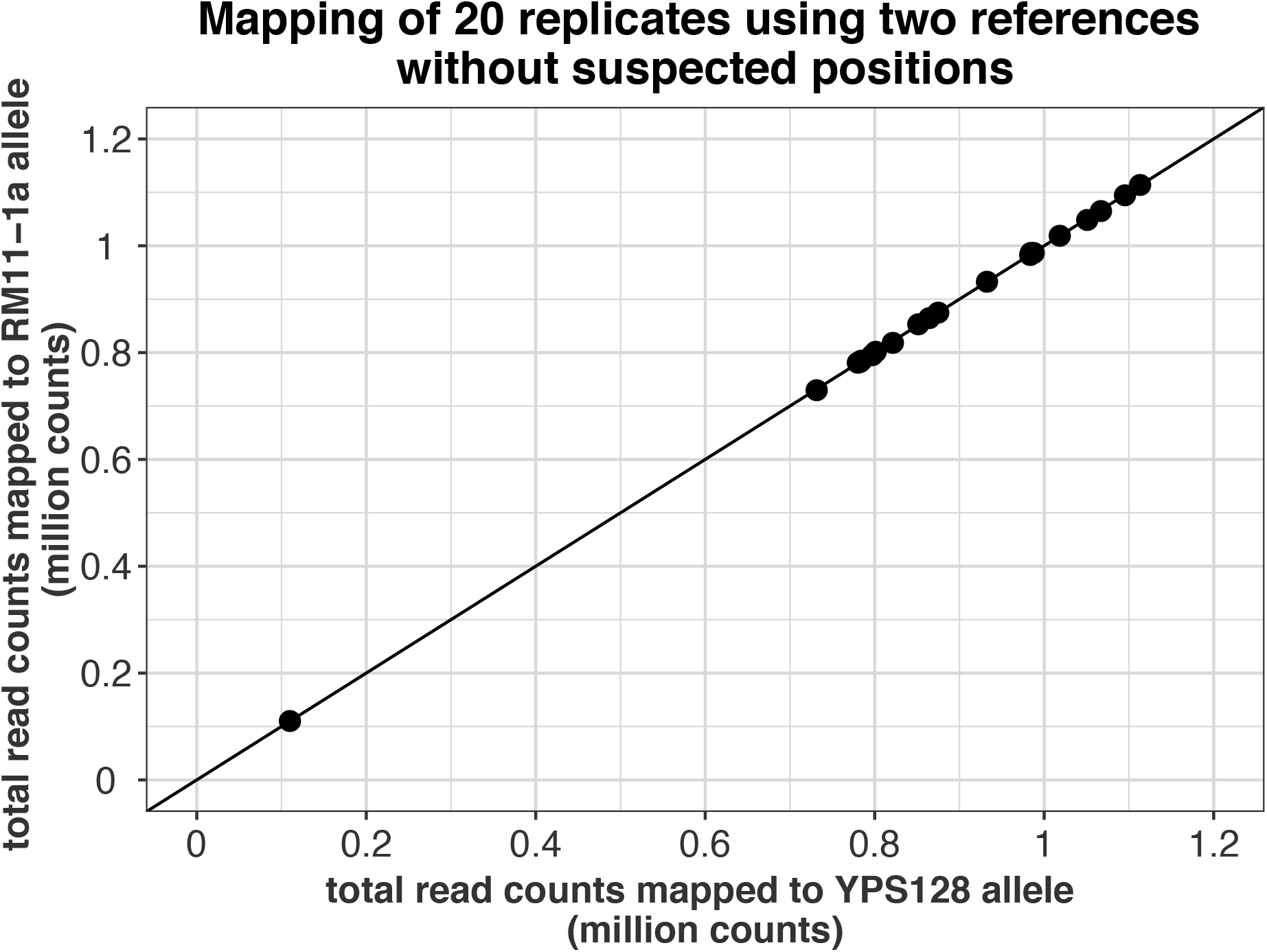
The mapping bias of DNA read counts are further eliminated by filtering out suspected variant positions.

**Figure S4:**
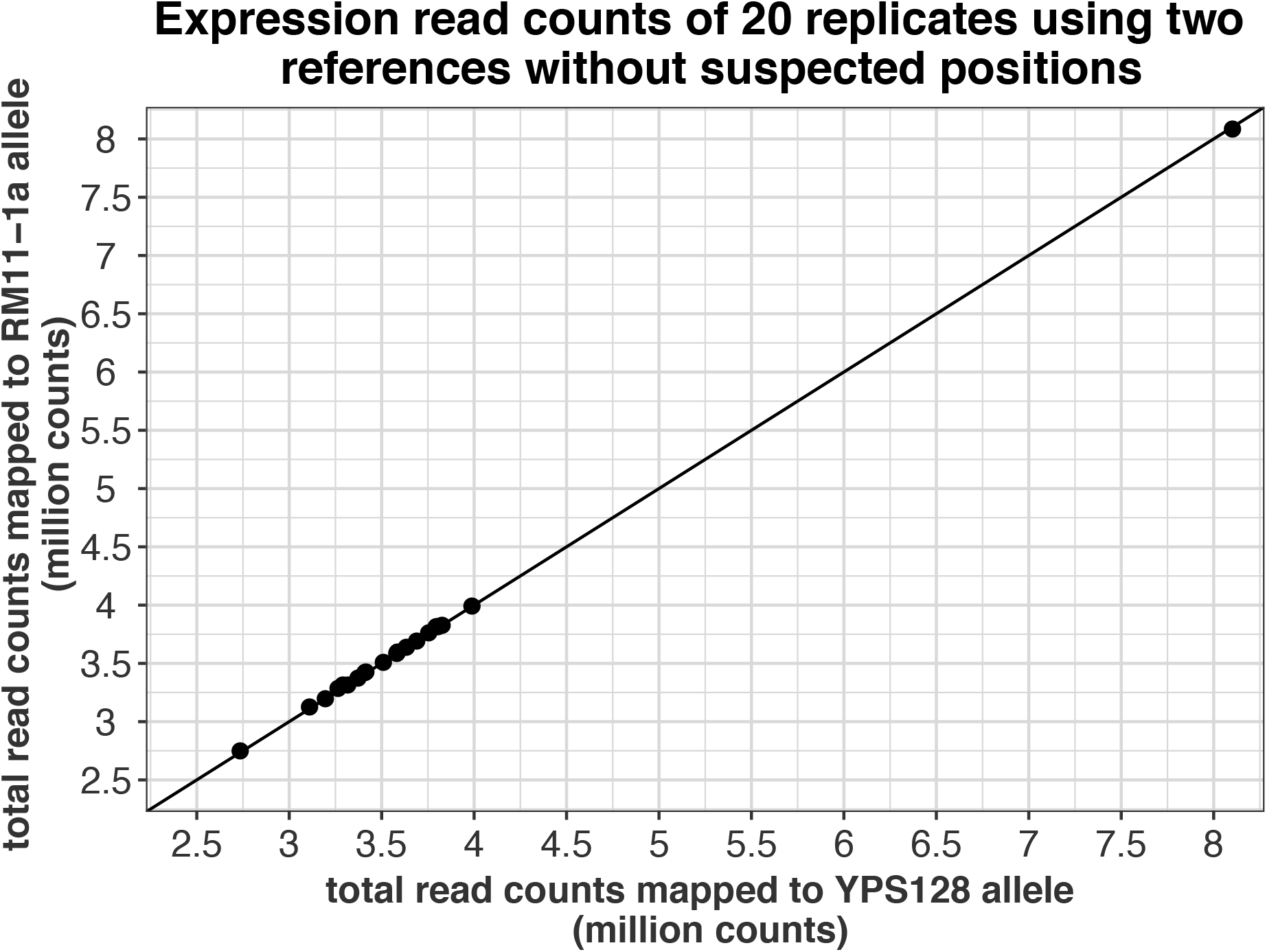
By using two references and removing all suspected variant positions, no mapping bias shown up for the RNA read counts.

**Figure S5:**
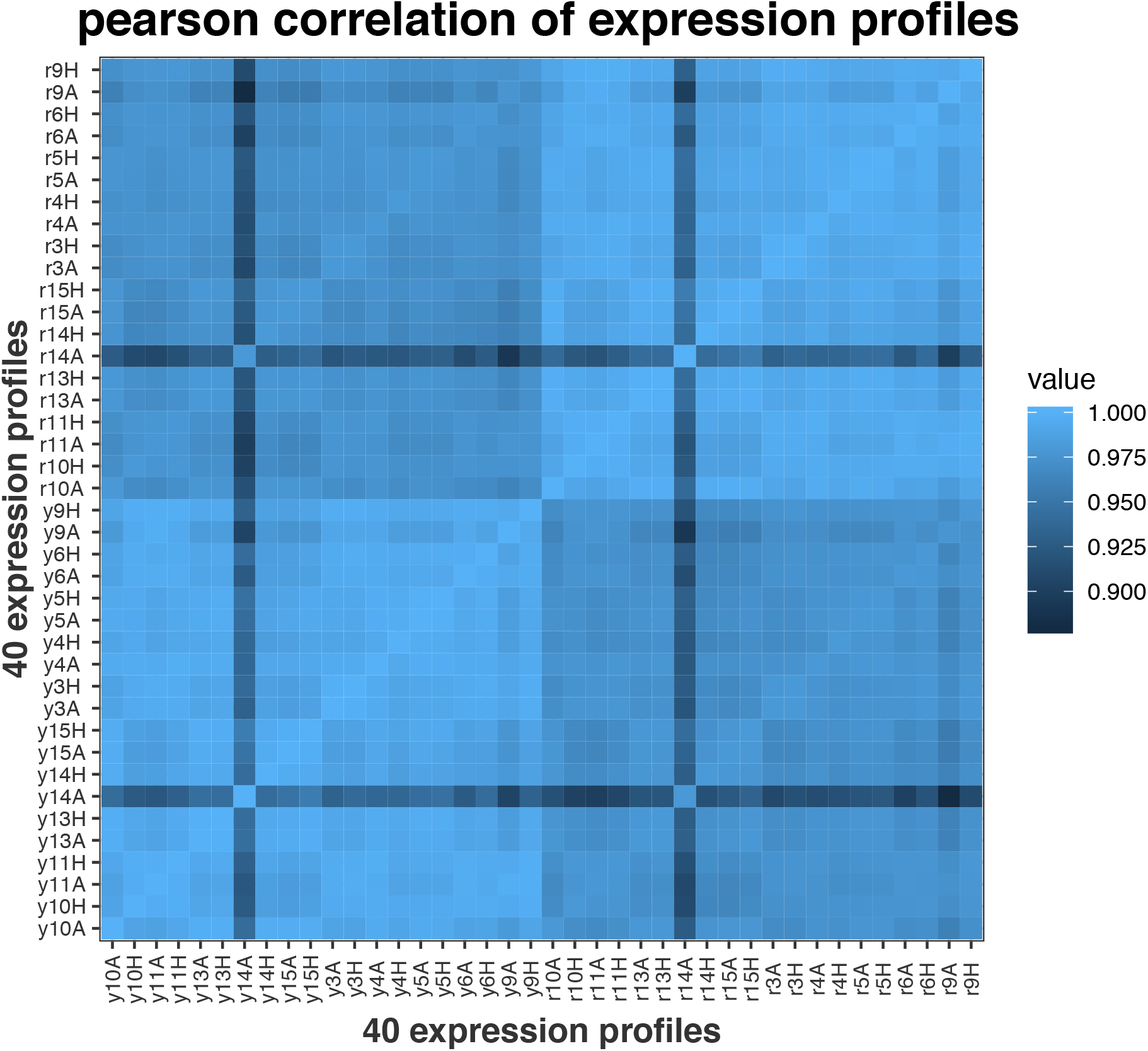
Correlation of 40 expression profiles. Profiles: r9A, y9A, r14A and y14A are removed for downstream analysis because of the low correlation with other replicates.

**Figure S6:**
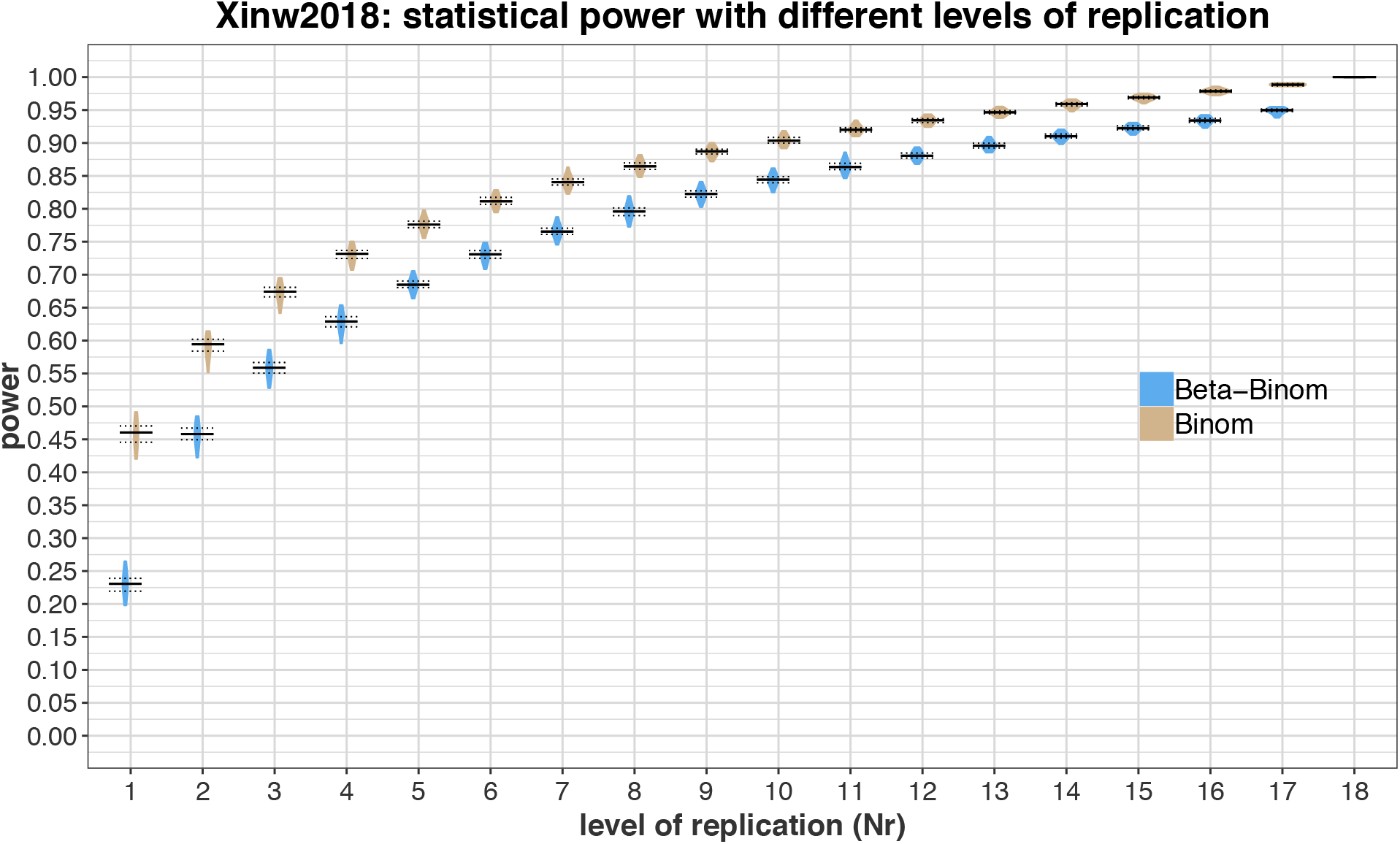
The statistical power of identifying significant cis-affected genes are plotted against increasing level of replication. The significant gene list from the best estimates (i.e. the betabinomial model with all 18 replicates) was used as a gold standard. The power increases for both models as more replicates used. Binomial model has higher statistical power than beta-binomial model.

## Acknowledgement

We thank A. Long, and TY. Wang for providing the yeast strain, T.Tsuboi for assistance in making yeast hybrid and culturing, J.Shan, R.Linder, J.Baldwin-Brown for assistance in making DNA and RNA libraries, M.Chakraborty for suggestions in assembling the genomes, J.Schraiber, A.Ramaiah and N.Zhao for thoughtful comments on the manuscript.

The work was supported by US National Institutes of Health (NIH) grant R01GM123303-1 (J.J.E.), University of California, Irvine setup funds (J.J.E). This work was made possible, in part, through access to the High Performance Computing Cluster of University of California, Irvine.

## References

Anders, Simon, Paul Theodor Pyl, and Wolfgang Huber. 2015. “HTSeq--a Python Framework to Work with High-Throughput Sequencing Data.” Bioinformatics 31 (2): 166–69.

Artieri, Carlo G., and Hunter B. Fraser. 2014. “Evolution at Two Levels of Gene Expression in Yeast.” Genome Research 24 (3): 411–21.

Bell, Graeme D. M., Nolan C. Kane, Loren H. Rieseberg, and Keith L. Adams. 2013. “RNA-Seq Analysis of Allele-Specific Expression, Hybrid Effects, and Regulatory Divergence in Hybrids Compared with Their Parents from Natural Populations.” Genome Biology and Evolution 5 (7): 1309–23.

Brem, Rachel B., Gaël Yvert, Rebecca Clinton, and Leonid Kruglyak. 2002. “Genetic Dissection of Transcriptional Regulation in Budding Yeast.” Science 296 (5568): 752–55.

Degner, Jacob F., John C. Marioni, Athma A. Pai, Joseph K. Pickrell, Everlyne Nkadori, Yoav Gilad, and Jonathan K. Pritchard. 2009. “Effect of Read-Mapping Biases on Detecting Allele-Specific Expression from RNA-Sequencing Data.” Bioinformatics 25 (24): 3207–12.

Emerson, J. J., Li-Ching Hsieh, Huang-Mo Sung, Tzi-Yuan Wang, Chih-Jen Huang, Henry Horng-Shing Lu, Mei-Yeh Jade Lu, Shu-Hsing Wu, and Wen-Hsiung Li. 2010. “Natural Selection on Cis and Trans Regulation in Yeasts.” Genome Research 20 (6): 826–36.

Emerson, J. J., and Wen-Hsiung Li. 2010. “The Genetic Basis of Evolutionary Change in Gene Expression Levels.” Philosophical Transactions of the Royal Society of London. Series B, Biological Sciences 365 (1552): 2581–90.

Garrison, Erik, and Gabor Marth. 2012. “Haplotype-Based Variant Detection from Short-Read Sequencing.” arXiv[q-bio.GN]. arXiv. http://arxiv.org/abs/1207.3907.

Gierliński, Marek, Christian Cole, Pietà Schofield, Nicholas J. Schurch, Alexander Sherstnev, Vijender Singh, Nicola Wrobel, et al. 2015. “Statistical Models for RNA-Seq Data Derived from a Two-Condition 48-Replicate Experiment.” Bioinformatics 31 (22): 3625–30.

Hodgins-Davis, Andrea, Daniel P. Rice, and Jeffrey P. Townsend. 2015. “Gene Expression Evolves under a House-of-Cards Model of Stabilizing Selection.” Molecular Biology and Evolution 32 (8): 2130–40.

Jacob, F., and J. Monod. 1961. “Genetic Regulatory Mechanisms in the Synthesis of Proteins.” Journal of Molecular Biology 3 (June): 318–56.

Kim, Daehwan, Geo Pertea, Cole Trapnell, Harold Pimentel, Ryan Kelley, and Steven L. Salzberg. 2013. “TopHat2: Accurate Alignment of Transcriptomes in the Presence of Insertions, Deletions and Gene Fusions.” Genome Biology 14 (4): R36.

King, M. C., and A. C. Wilson. 1975. “Evolution at Two Levels in Humans and Chimpanzees.” Science 188 (4184): 107–16.

Koren, Sergey, Arang Rhie, Brian P. Walenz, Alexander T. Dilthey, Derek M. Bickhart, Sarah B. Kingan, Stefan Hiendleder, John L. Williams, Timothy P. L. Smith, and Adam M. Phillippy. 2018. “Complete Assembly of Parental Haplotypes with Trio Binning.” bioRxiv. https://doi.org/10.1101/271486.

Koren, Sergey, Brian P. Walenz, Konstantin Berlin, Jason R. Miller, Nicholas H. Bergman, and Adam M. Phillippy. 2017. “Canu: Scalable and Accurate Long-Read Assembly via Adaptive K-Mer Weighting and Repeat Separation.” Genome Research 27 (5): 722–36.

Kurtz, Stefan, Adam Phillippy, Arthur L. Delcher, Michael Smoot, Martin Shumway, Corina Antonescu, and Steven L. Salzberg. 2004. “Versatile and Open Software for Comparing Large Genomes.” Genome Biology 5 (2): R12.

Lam, Ka-Kit, Kurt LaButti, Asif Khalak, and David Tse. 2015. “FinisherSC: A Repeat-Aware Tool for Upgrading de Novo Assembly Using Long Reads.” Bioinformatics 31 (19): 3207–9.

Langmead, Ben, and Steven L. Salzberg. 2012. “Fast Gapped-Read Alignment with Bowtie 2.” Nature Methods 9 (4): 357–59.

Li, Heng, and Richard Durbin. 2009. “Fast and Accurate Short Read Alignment with Burrows-Wheeler Transform.” Bioinformatics 25 (14): 1754–60.

Li, Heng, Bob Handsaker, Alec Wysoker, Tim Fennell, Jue Ruan, Nils Homer, Gabor Marth, Goncalo Abecasis, Richard Durbin, and 1000 Genome Project Data Processing Subgroup. 2009. “The Sequence Alignment/Map Format and SAMtools.” Bioinformatics 25 (16): 2078–79.

Mack, Katya L., Polly Campbell, and Michael W. Nachman. 2016. “Gene Regulation and Speciation in House Mice.” Genome Research 26 (4): 451–61.

Mcclintock, B. 1956. “Controlling Elements and the Gene.” Cold Spring Harbor Symposia on Quantitative Biology 21: 197–216.

McManus, C. Joel, Joseph D. Coolon, Michael O. Duff, Jodi Eipper-Mains, Brenton R. Graveley, and Patricia J. Wittkopp. 2010. “Regulatory Divergence in Drosophila Revealed by mRNA-Seq.” Genome Research 20 (6): 816–25.

McManus, C. Joel, Gemma E. May, Pieter Spealman, and Alan Shteyman. 2014. “Ribosome Profiling Reveals Post-Transcriptional Buffering of Divergent Gene Expression in Yeast.” Genome Research 24 (3): 422–30.

Metzger, Brian P. H., Patricia J. Wittkopp, and Joseph D. Coolon. 2017. “Evolutionary Dynamics of Regulatory Changes Underlying Gene Expression Divergence among Saccharomyces Species.” Genome Biology and Evolution 9 (4): 843–54.

Ohno, S. 1972. “So Much ‘Junk’ DNA in Our Genome.” Brookhaven Symposia in Biology 23: 366–70.

Picelli, Simone, Omid R. Faridani, Asa K. Björklund, Gösta Winberg, Sven Sagasser, and Rickard Sandberg. 2014. “Full-Length RNA-Seq from Single Cells Using Smart-seq2.” Nature Protocols 9 (1): 171–81.

Renaud, Gabriel, Udo Stenzel, Tomislav Maricic, Victor Wiebe, and Janet Kelso. 2015. “deML: Robust Demultiplexing of Illumina Sequences Using a Likelihood-Based Approach.” Bioinformatics 31 (5): 770–72.

R Foundation for Statistical Computing, Vienna, Austria. n.d. “R Core Team (2018). R: A Language and Environment for Statistical Computing.” https://www.r-project.org/.

Rhoné, Bénédicte, Cédric Mariac, Marie Couderc, Cécile Berthouly-Salazar, Issaka Salia Ousseini, and Yves Vigouroux. 2017. “No Excess of Cis-Regulatory Variation Associated with Intraspecific Selection in Wild Pearl Millet (Cenchrus Americanus).” Genome Biology and Evolution 9 (2): 388–97.

Robinson, Mark D., and Gordon K. Smyth. 2007. “Moderated Statistical Tests for Assessing Differences in Tag Abundance.” Bioinformatics 23 (21): 2881–87.

Robinson, Mark D., and Gordon K. Smyth. 2008. “Small-Sample Estimation of Negative Binomial Dispersion, with Applications to SAGE Data.” Biostatistics 9 (2): 321–32.

Romero, Irene Gallego, Ilya Ruvinsky, and Yoav Gilad. 2012. “Comparative Studies of Gene Expression and the Evolution of Gene Regulation.” Nature Reviews. Genetics 13 (7): 50516.

Schaefke, Bernhard, J. J. Emerson, Tzi-Yuan Wang, Mei-Yeh Jade Lu, Li-Ching Hsieh, and Wen-Hsiung Li. 2013. “Inheritance of Gene Expression Level and Selective Constraints on Trans- and Cis-Regulatory Changes in Yeast.” Molecular Biology and Evolution 30 (9): 2121–33.

Schurch, Nicholas J., Pietá Schofield, Marek Gierliński, Christian Cole, Alexander Sherstnev, Vijender Singh, Nicola Wrobel, et al. 2016. “How Many Biological Replicates Are Needed in an RNA-Seq Experiment and Which Differential Expression Tool Should You Use?” RNA 22 (6): 839–51.

Signor, Sarah A., and Sergey V. Nuzhdin. 2018. “The Evolution of Gene Expression in Cis and Trans.” Trends in Genetics: TIG 34 (7): 532–44.

Simão, Felipe A., Robert M. Waterhouse, Panagiotis Ioannidis, Evgenia V. Kriventseva, and Evgeny M. Zdobnov. 2015. “BUSCO: Assessing Genome Assembly and Annotation Completeness with Single-Copy Orthologs.” Bioinformatics 31 (19): 3210–12.

Sniegowski, Paul D., Peter G. Dombrowski, and Ethan Fingerman. 2002. “Saccharomyces Cerevisiae and Saccharomyces Paradoxus Coexist in a Natural Woodland Site in North America and Display Different Levels of Reproductive Isolation from European Conspecifics.” FEMS Yeast Research 1 (4): 299–306.

Tirosh, Itay, Sharon Reikhav, Avraham A. Levy, and Naama Barkai. 2009. “A Yeast Hybrid Provides Insight into the Evolution of Gene Expression Regulation.” Science 324 (5927): 659–62.

Walker, Bruce J., Thomas Abeel, Terrance Shea, Margaret Priest, Amr Abouelliel, Sharadha Sakthikumar, Christina A. Cuomo, et al. 2014. “Pilon: An Integrated Tool for Comprehensive Microbial Variant Detection and Genome Assembly Improvement.” PloS One 9 (11): e112963.

Wittkopp, Patricia J., Belinda K. Haerum, and Andrew G. Clark. 2008. “Regulatory Changes Underlying Expression Differences within and between Drosophila Species.” Nature Genetics 40 (3): 346–50.

Wray, Gregory A. 2007. “The Evolutionary Significance of Cis-Regulatory Mutations.” Nature Reviews. Genetics 8 (3): 206–16.

Yue, Jia-Xing, Jing Li, Louise Aigrain, Johan Hallin, Karl Persson, Karen Oliver, Anders Bergström, et al. 2017. “Contrasting Evolutionary Genome Dynamics between Domesticated and Wild Yeasts.” Nature Genetics 49 (6): 913–24.

Zhao, Hao, Zhifu Sun, Jing Wang, Haojie Huang, Jean-Pierre Kocher, and Liguo Wang. 2014. “CrossMap: A Versatile Tool for Coordinate Conversion between Genome Assemblies.” Bioinformatics 30 (7): 1006–7.

